# A hollow fiber membrane-based liver organoid-on-a-chip model for examining drug metabolism and transport

**DOI:** 10.1101/2024.08.12.607504

**Authors:** Adam Myszczyszyn, Anna Münch, Vivian Lehmann, Theo Sinnige, Frank G. van Steenbeek, Manon Bouwmeester, Roos-Anne Samsom, Marit Keuper-Navis, Thomas K. van der Made, Daniel Kogan, Sarah Braem, Luc J. W. van der Laan, Hossein Eslami Amirabadi, Evita van de Steeg, Rosalinde Masereeuw, Bart Spee

**Affiliations:** Department of Clinical Sciences, Faculty of Veterinary Medicine, Utrecht University, Utrecht, The Netherlands; Institute for Risk Assessment Sciences (IRAS), Utrecht University, Utrecht, The Netherlands; Department of Metabolic Health Research, Netherlands Organisation for Applied Scientific Research (TNO), Leiden, The Netherlands; Division of Pharmacology, Department of Pharmaceutical Sciences, Faculty of Science, Utrecht University, Utrecht, The Netherlands; Department of Surgery, Erasmus MC Transplant Institute, University Medical Center Rotterdam, Rotterdam, The Netherlands; AZAR Innovations, Utrecht, The Netherlands

## Abstract

Liver-on-a-chip models predictive for both metabolism as well as canalicular and blood transport of drug candidates in humans are lacking. Here, we established an advanced, bioengineered and animal component-free hepatocyte-like millifluidic system based on 3D hollow fiber membranes (HFMs), recombinant human laminin 332 coating and adult human stem cell-derived organoids. Organoid fragments formed polarized and tight monolayers on HFMs with improved hepatocyte-like maturation, as compared to standard 3D organoid cultures in Matrigel from matched donors. Gene expression profiling and immunofluorescence revealed that hepatocyte-like monolayers expressed a broad panel of phase I (e.g., CYP3A4, CYP2D6) and II (UGTs, SULTs) drug-metabolizing enzymes and drug transporters (e.g., OATP1B3, MDR1 and MRP3). Moreover, statically cultured monolayers displayed phase I and II metabolism of a cocktail of six relevant compounds, including midazolam and 7-hydroxycoumarin. We also demonstrated the disposition of midazolam in the basal/blood-like circulation and apical/canalicular-like compartment of the millifluidic chip. Finally, we connected the system to the other two PK/ADME-most relevant organ systems, *i.e.* small intestine- and kidney proximal tubule-like to study the bioavailability of midazolam and coumarin, and excretion of metformin. In conclusion, we generated a proof-of-concept liver organoid-on-a-chip model for examining metabolism and transport of drugs, which can be further developed to predict PK/ADME profiles in humans.

## Introduction

Rodent models used during preclinical development are poorly predictive for behaviors of drug candidates in humans,^[1]^ and the lack of translation results in 90% failure of those compounds during clinical development^[2]^. This problem has recently been acknowledged by the US Food and Drug Administration (FDA) Modernization Act 2.0 of 2022 that allows for utilizing relevant *in vitro* models instead of animal testing for every new non-clinical drug development protocol.^[3]^ The lack of predictiveness for pharmacokinetics’ (PK) parameters in humans, *i.e.* absorption, distribution, metabolism and excretion (ADME), is the second major biological reason for high attrition rates of drug candidates in phase I of clinical development, after safety.^[1,4]^ This underscores the need for more predictive *in vitro* preclinical systems for major PK/ADME-related organs such as the liver.

In the past years, advanced bioengineered organ-on-a-chip (microphysiological) technologies have emerged as most promising *in vitro* models for predicting human PK/ADME profiles, as they can closely recapitulate the architecture, environmental factors and functions of tissues of origin.^[5–7]^ Based on commercially manufactured chips (from e.g., *Emulate, Inc.*; *Javelin Biotech, Inc.*; *CN Bio Innovations Ltd.* and *TissUse GmbH*), these micro- and millifluidic technologies improved the functionality of primary human hepatocytes (PHHs) over a longer period of culture by recapitulating their native canalicular polarization thanks to extracellular matrix (ECM) sandwich-culture or 3D/spheroid growth as well as by incorporating supporting cells and flow-induced mechanical forces.^[8–11]^ However, this makes PHHs-on-a-chip systems only suitable for studying drug uptake and metabolism, as the apical/canalicular-like compartment between cells is inaccessible for examining passive diffusion (transcellular and paracellular) and active transport of drugs and their metabolites^[12]^. Thus, liver-on-a-chip models with an apical/canalicular-like compartment that can be directly sampled to capture biliary excretion and enterohepatic circulation of drug candidates are an unmet need.^[5,7]^ Previously, human embryonic stem cell (ESC)- and induced pluripotent stem cell (iPSC)-derived hepatocyte-like monolayers on Transwell membranes displayed columnar epithelial polarization with an accessible apical/canalicular- like compartment. Though, perfusion was not applied, and these hepatocyte-like cells were still immature, as they exhibited transcriptomic profiles similar to the embryonic liver.^[13]^

Hollow fiber membranes (HFMs) have emerged as perfusable scaffolds that are suitable for generating polarized epithelial monolayers with two accessible micro-/millifluidic compartments. These 3D tubular-shaped and porous membranes are made of a biocompatible, stable and low-cost material such as MicroPES (polyethersulfone).^[14]^ In contrast to flat 2D Transwell membranes, HFMs were shown to enhance cell differentiation^[15]^, possibly thanks to 3D geometrical properties^[16]^. Previously, we took advantage of HFMs to establish kidney proximal tubule-^[17]^ and small intestine-on-a-chip^[14]^ systems based on immortalized human cell lines. In addition, we created mouse^[18]^ and human^[19]^ bile duct-on-a-chip HFM-based models using liver organoids. However, an organoid-derived HFM-based liver-on-a-chip system has not been reported to date.

Human stem cell-derived 3D organoids from the healthy adult liver, so-called intrahepatic cholangiocyte organoids (ICOs)^[20]^, are a robust cell source for establishing liver-on-a-chip systems. ICOs can be expanded in Matrigel droplet-based cultures in expansion medium via serial passaging over a long-term while retaining genetic stability and potential for biobanking. They are enriched in LGR5-positive bipotent progenitor cells that can maturate into hepatocyte-like cells upon a switch to a differentiation medium. These cells maintain major functional features of individuals, including variations in the expression of metabolizing enzymes and transporters of drugs.^[21]^

Current liver-on-a-chip models do not meet the 3Rs’ principles^[3]^, as they utilize fetal bovine serum-rich media and animal-derived extracellular matrix (ECM) components such as Matrigel from mouse sarcoma, rat tail Collagen I and bovine Fibronectin^[9]^. These components are undefined, which results in experimental variability, poor reproducibility and the lack of standardization of liver-on-a-chip systems, potentially blocking their approval by regulatory agencies.^[3]^ Alternative human tissue extract-derived ECM components like Collagen IV from placenta^[9]^ are not fully defined and are also quite costly. In contrast, recombinant human laminins such as laminin 111, 332 and 511 can aid animal component- free standardization and are more cost effective. Moreover, these laminins are expressed in the quiescent^[22]^ and regenerating^[23]^ adult human liver, and were shown to maintain the functionality of 2D monolayer cultures of PHHs^[22]^ as well as to drive hepatocyte-like differentiation of ESCs^[24,25]^ and iPSCs^[26–28]^. Previously, we demonstrated that a chemically defined synthetic hydrogel, PIC, and laminin 111 can replace Matrigel in culturing ICOs.^[29,30]^ Together, recombinant human laminins can offer a biologically relevant alternative for ECM components in liver-on-a-chip systems.

Animal component-free liver-on-a-chip models that display improved maturation and tight epithelial barriers with apical-to-basolateral columnar polarization are necessary for a successful human *in vitro*-to-*in vivo* (IVIVE) extrapolation of PK/ADME profiles of drug candidates. Here, we generated an advanced, bioengineered hepatocyte-like millifluidic system based on HFMs, recombinant human laminin 332 coating, human liver organoid fragments and serum-free media with defined growth factors. This system is metabolically active and allows for the sampling of the basal and apical compartment to examine transport of parent compounds and their metabolites across polarized cell monolayers. Thus, our proof-of-concept model has application potential in determining experimental PK/ADME kinetic values of drug candidates.

## Materials and Methods

### Liver organoids

The use of tissue from healthy adult liver transplant recipients at the Erasmus Medical Center Rotterdam (EMC, Netherlands) for research purposes was ethically approved (MEC-2014- 060). These included: donor 1 (male, middle-aged), donor 2 (female, middle-aged), donor 3 (male, middle-aged), donor 4 (female, middle-aged) and donor 5 (female, middle-aged). ICOs were established, as described previously^[21,31]^. ICOs were cultured in 25-µl droplets of Growth Factor Reduced Matrigel (Corning), 70% (v/v; in expansion medium, EM) and 100% for batches ≥ 9 mg/mL and ≤ 9 mg/mL of proteins, respectively. Matrigel droplets with ICO fragments were cultured in EM that contained advanced DMEM/F12 (Thermo Fisher Scientific), 1% (v/v) penicillin-streptomycin (Thermo Fisher Scientific), 1% (v/v) GlutaMAX (Thermo Fisher Scientific), 10 mM HEPES (Thermo Fisher Scientific), 2% (v/v) B27 supplement without vitamin A (serum-free, Thermo Fisher Scientific), 1% (v/v) N2 supplement (serum-free, Thermo Fisher Scientific), 1.25 mM N-acetylcysteine (Sigma- Aldrich), 10 mM nicotinamide (Sigma-Aldrich), 2% (v/v) human R-spondin-3-Fc Fusion Protein Conditioned Medium (ImmunoPrecise Antibodies), 50 ng/mL recombinant human EGF (Peprotech), 100 ng/mL recombinant human FGF10 (Peprotech), 25 ng/mL recombinant human HGF (Peprotech), 10 nM human gastrin (Leu15 analog, AnaSpec), 5 µM A83-01 (STEMCELL Technologies) and 10 µM forskolin (Sigma-Aldrich). Cultures were incubated in a humidified atmosphere at 37°C and 5% CO_2_. EM was refreshed every 2-3 days. Cultures were passaged every 4-7 days (until the confluency was reached) at a 1:3-1:5 split ratio by mechanical fragmentation. Passages 4-12. were used for experiments. To induce hepatocyte-like maturation, ICOs expanded for 4-7 days were further cultured for 3 days in pre-differentiation medium (pre-DM), *i.e.* EM supplemented with 25 ng/mL recombinant human BMP7 (Peprotech), and for 7 days in differentiation medium (DM) consisting of advanced DMEM/F12 (Thermo Fisher Scientific), 1% (v/v) penicillin- streptomycin (Thermo Fisher Scientific), 1% (v/v) GlutaMAX (Thermo Fisher Scientific), 10 mM HEPES (Thermo Fisher Scientific), 2% (v/v) B27 supplement without vitamin A (serum- free, Thermo Fisher Scientific), 1% (v/v) N2 supplement (serum-free, Thermo Fisher Scientific), 1.25 mM N-acetylcysteine (Sigma-Aldrich), 50 ng/mL recombinant human EGF (Peprotech), 100 ng/mL recombinant human FGF19 (Peprotech), 25 ng/mL recombinant human HGF (Peprotech), 25 ng/mL recombinant human BMP7 (Peprotech), 10 nM human gastrin (Leu15 analog, AnaSpec), 0.5 µM A83-01 (STEMCELL Technologies), 10 µM DAPT (Selleck Chemicals) and 30 µM dexamethasone (Sigma-Aldrich). For gene expression profiling, ICOs were cultured in DM for 10 days without the pre-DM phase. DM was refreshed every 2-3 days. Brightfield images were acquired with an EVOS M5000 Imaging System (Thermo Fisher Scientific).

### A hepatocyte-like HFM-based system

MicroPES (polyethersulfone) TF10/PVP HFMs with an internal diameter of 300 µm, wall thickness of 100 µm and a maximum pore size of 0.5 µm were extracted from a SepaPlas 06 plasma filter (MEDICA) and cut into 3- or 4-cm pieces for static and chip assays, respectively. HFM pieces were then sterilized in 70% (v/v) EtOH for 30 min on a roller and washed 3x with sterile 1x PBS. L-DOPA (3,4-dihydroxy-L-phenylalanine, Sigma-Aldrich) was dissolved to 2 mg/mL in 10 mM Tris-HCl buffer (pH 8.5) on a carousel (at 10 rpm) at 37°C overnight. The solution was filter-sterilized and added to sterile HFM pieces that were then incubated on a carousel (at 10 rpm) at 37°C for four hrs. Next, the solution was aspirated, and L-DOPA-coated HFM pieces were washed 3x with sterile 1x PBS and stored at 4°C for one week or used immediately for ECM coating. For static assays, Growth Factor Reduced Matrigel (Corning), or recombinant human laminin 111, 211, 332, 411 or 511 (BioLamina) were diluted with sterile 1x PBS to 10 µg/mL (according to the manufacturer’s recommendation) in the volume of 5 mL in 5-mL Protein LoBind tubes (Eppendorf). For chip assays, laminin 332 was used exclusively. Up to 15 HFM pieces were transferred to an ECM solution and incubated on a carousel (10 rpm) at 37°C for two hrs. ECM-coated HFM pieces were immediately used for seeding ICO fragments. Matrigel droplets with ICOs were mechanically harvested using advanced DMEM/F12 (Thermo Fisher Scientific) with 1% (v/v) penicillin-streptomycin (Thermo Fisher Scientific), 1% (v/v) GlutaMAX (Thermo Fisher Scientific) and 10 mM HEPES (Thermo Fisher Scientific), and Matrigel was removed by repeated pipetting and washings. ICO pellets were mechanically fragmented by extensive pipetting in 1 mL of the latter medium. For static assays, HFM pieces were seeded with ICO fragments in 2-mL tubes at a ratio of two pieces per 1 mL of EM with ICO fragments harvested from eight confluent Matrigel droplets. For chip assays, single HFM pieces were placed in an in-house 3D-printed holder using a biocompatible resin and fixed with dental glue (Gi-Mask Automix, Coltène). The holder was designed in SolidWorks (Dessault Systèmes) and stereolitography-printed using a RapidShape S60 LED DLP printer (Rapid Shape). The design is the subject of a patent process. HFM-carrying holders were seeded with ICO fragments in 5-mL tubes at a ratio of one holder per 3.5 mL of EM with ICO fragments harvested from 28 confluent Matrigel droplets. For static assays, tubes were incubated horizontally in a humidified atmosphere at 37°C and 5% CO_2_ with a gentle resuspension by a 360° rotation every 45 min (4x), totalling four hrs of incubation. Next, cell- laden HFMs were transferred to 6-well plates with 2 mL of EM (one HFM per well) and cultured for 4-7 days in a humidified atmosphere at 37°C and 5% CO_2_ without medium change, until the confluency was reached. For chip assays, tubes were incubated vertically on a carousel (at < 0.5 rpm) at 37°C for four hrs. Cell-laden HFMs in holders were transferred to 6-well plates with 6 mL of EM (one holder per well) and cultured for 4-7 days in a humidified atmosphere at 37°C and 5% CO_2_, without medium change, until the confluency was reached. To induce hepatocyte-like maturation, HFM-expanded monolayers were further statically cultured for 3 days in pre-DM and for 7 or 14 or 21 days in DM. HFM-expanded monolayers in holders were further statically cultured for 3 days in pre-DM and for 7 days in DM. Brightfield images were acquired with an EVOS M5000 Imaging System (Thermo Fisher Scientific).

### Real-time qPCR

Matrigel droplets with ICOs (eight per condition) were harvested, and Matrigel was removed by incubating with Cell Recovery Solution (Corning), according to the manufacturer’s instructions. Total RNA was isolated from HFM-grown monolayers (six per condition), ICOs and liver tissue specimens using the RNeasy Mini Kit (Qiagen), according to the manufacturer’s instructions, including supplementing RLT buffer with β-mercaptoethanol and treatment of lysates with DNase I. RNA was quantified using a D-1000 NanoDrop spectrophotometer (Thermo Fisher Scientific). 1 µg RNA was reverse-transcribed into cDNA using the iScript cDNA synthesis kit (Bio-Rad), according to the manufacturer’s instructions. Synthesized cDNA was diluted 7x. Relative mRNA of genes of interest was quantified by real-time qPCR reactions performed in a CFX384 Touch Real-Time PCR Detection System (thermal cycler, Bio-Rad). The iQ SYBR Green Supermix (Bio-Rad) and 0.7 µM primers were used (Table 1) that were designed in-house and synthesized at Eurogentec, and then validated by performing a temperature gradient analysis. A 2-step reaction was performed at 60°C, *i.e.* combined annealing and elongation, with 39 cycles. Primer specificity was tested by melting curve analysis and running reactions with a negative control (without cDNA). Primer efficiency was tested by running standard curves with pooled cDNA samples. Relative mRNA expression values were normalized to the averaged endogenous controls *RPL19* (structural protein) and *HPRT* (metabolic enzyme) using the 2^-ΔΔCt^ method. A cut-off C_q_ value was 35.

**Table 1.**
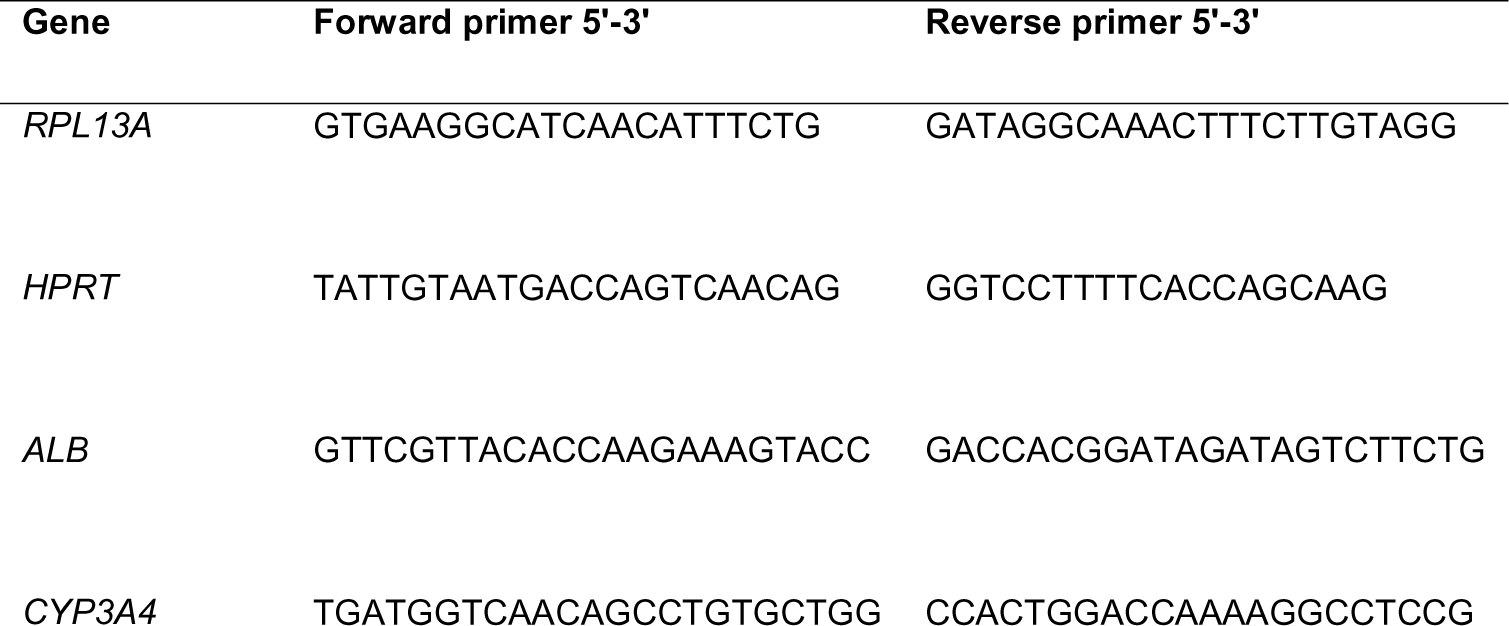
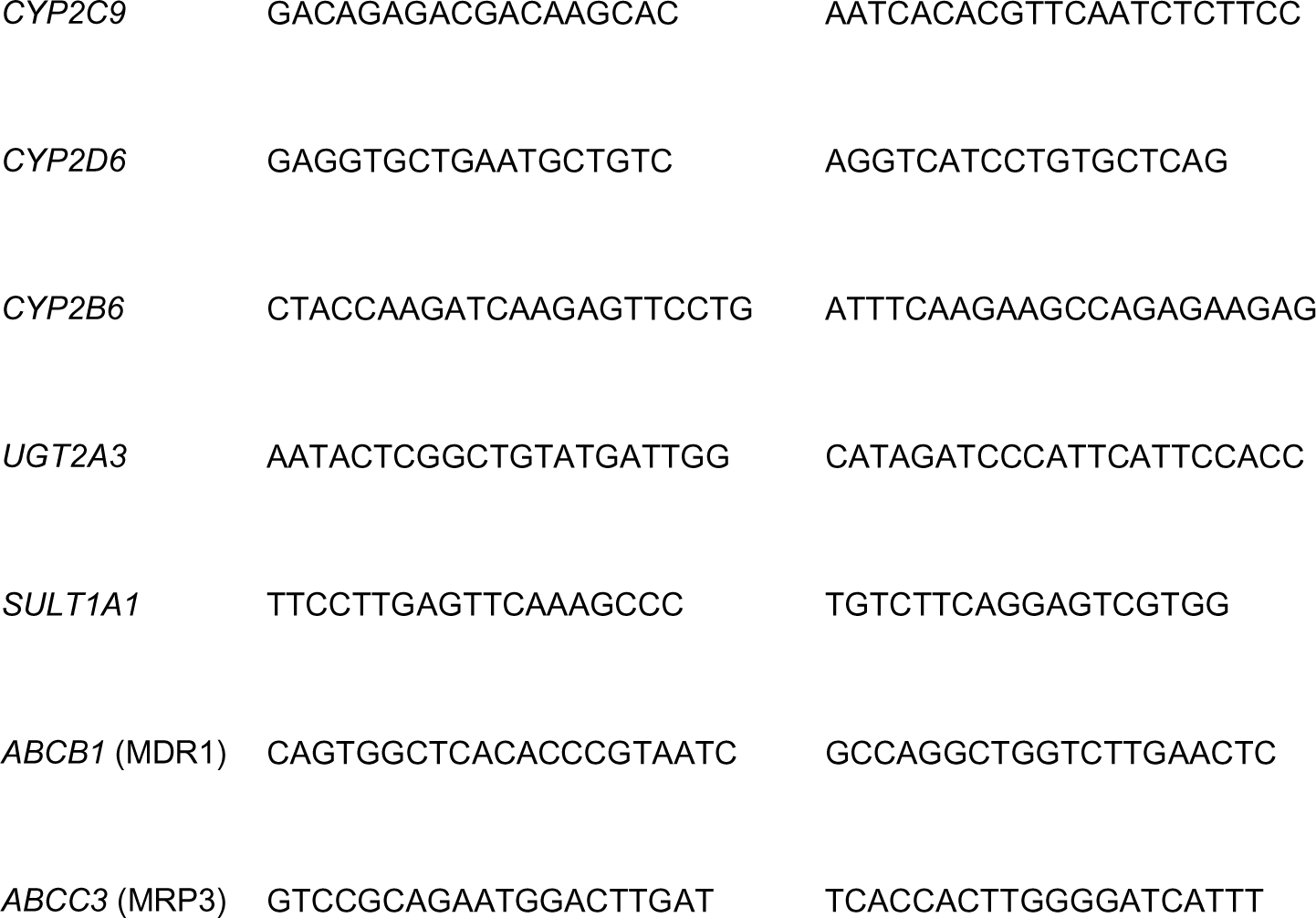
Primer sequences for real-time qPCR.

### Bulk mRNA sequencing

Matrigel droplets with ICOs (eight per condition) were mechanically harvested, and Matrigel was removed by incubating with Cell Recovery Solution (Corning), according to the manufacturer’s instructions. Total RNA was isolated from HFM-grown monolayers (six per condition), ICOs and liver tissue specimens using the RNeasy Mini Kit (Qiagen), according to the manufacturer’s instructions, including supplementing RLT buffer with β-mercaptoethanol and treatment of lysates with DNase I. The amount and quality of RNA was measured with the Agilent RNA 6000 Nano Kit and an Agilent 2100 Bioanalyzer (Agilent Technologies), according to the manufacturer’s instructions. All samples had a RNA integrity number (RIN) value of ≥ 8. A minimum of 50 µl RNA at a minimum concentration of 4 ng/µl (200 ng in total) was submitted for mRNA library preparation (Truseq RNA stranded polyA enrichment) and sequencing at the Utrecht Sequencing Facility, according to internal protocols. The libraries were sequenced using the NextSeq2000 platform (Illumina), producing 1 x 50 bp reads, ∼20M reads per sample, 1,000M reads per flow cell (P3, 50 cycles). ICO and liver tissue samples were sequenced separately from monolayer samples. mRNA reads were aligned to the human reference genome GRCh37 using the STAR software version 2.4.2a (https://github.com/UMCUGenetics/RNASeq-NF/). Datasets were corrected for batch effects using Combat_seq^[32]^ and normalized for library size using trimmed mean of M values (TMM)^[33]^. Fold changes were calculated in edgeR^[34]^. For comparisons of gene expression, the exact test was used to calculate P-values that were corrected for the false discovery rate (FDR) of 5% using the Benjamini-Hochberg method. Genes were considered differentially expressed at adjusted P-value <L0.05. The principal component analysis plot and heatmaps of differentially expressed genes were created using ggplot2^[35]^. For the functional annotation, The Database for Annotation, Visualization and Integrated Discovery (DAVID) was used (https://david.ncifcrf.gov/). Differentially expressed genes were subjected to a cut-off for both adjusted P-value < 0.05 and log2 fold change ≥ 2 (actual fold change ≥ 4) for monolayers DM 7 days over organoids DM 10 days (upregulation in monolayers). 383 genes were selected. Using the entire human (Homo sapiens) genome as a background for the ENSEMBL_GENE_ID identifier, 350 genes (IDs) were identified and analyzed. Default parameters were used, except for the Enrichment Thresholds (EASE Scores, modified Fisher-exact P-values) and adjusted P-values (Benjamini-Hochberg correction) that were subjected to the cut-off of < 0.05 for all terms in each of 59 clusters. By this, 18 clusters were determined.

### Immunofluorescence microscopy

For whole-mount immunofluorescence, HFM-grown monolayers were washed once in 1x PBS, fixed in 4% (v/v) paraformaldehyde (PFA) on a roller at 4°C for 45 min. Next, PFA was removed and monolayers were washed 3x with 1x PBS. Monolayers were cut into 1-cm (or 1.5-cm) pieces and blocked on a rocker at room temperature (RT) for one h in a dilution/washing buffer containing 1x PBS, 1% (w/v) BSA and 0.1% (v/v) Tween 20 that was supplemented with 10% (v/v) goat serum (Sigma-Aldrich). Monolayer pieces were then incubated with primary antibodies in the buffer on a rocker in the darkness at 4°C overnight. These included rabbit anti-ZO1 (1:50, Thermo Fisher Scientific, 40-2300), rabbit anti-OATP1B3 (1:50, Novus Biologicals, NBP1-80980), rabbit anti-OCT1 (1:50, Novus Biologicals, NBP1-59464), mouse anti-MDR1 (1:50, Santa Cruz Biotechnology, sc-55510), rabbit anti-MRP2 (1:50, Abcam, ab187644), rabbit anti-MRP3 (1:50, Novus Biologicals, NBP2-37923) and phalloidin (a toxin against F-actin) conjugated to Alexa Fluor 488 (1:400, Thermo Fisher Scientific, A12379). Next, monolayer pieces were washed in the buffer 3x for 20 min on a rocker in the darkness. Incubation with secondary antibodies in the buffer, *i.e.* goat anti-rabbit-Alexa Fluor 488 (1:100, Thermo Fisher Scientific, A11008) and goat anti- mouse-Alexa Fluor 488 (1:100, Thermo Fisher Scientific, A11029), and with DAPI (1 μg/mL, Thermo Fisher Scientific) was performed for two hrs on a rocker in the darkness at RT. Then, monolayer pieces were washed in the buffer 3x for 10 min on a rocker in the darkness, followed by mounting on Lab-Tek II Chambered Coverglass (Nunc) with ProLong Diamond Antifade Mountant (Thermo Fisher Scientific) and immediate imaging. For immunofluorescence of paraffin sections, confluent HFM-grown monolayers were washed once in 1x PBS, fixed in 4% (v/v) PFA on a roller at 4°C for 45 min. Next, PFA was removed and monolayers were washed 3x in 1x PBS. Monolayers were cut into 0.5-cm pieces and mounted vertically in 2.5% (w/v) agarose droplets. Agarose droplets were dehydrated overnight in 70% (v/v) EtOH, 96% (v/v) EtOH, 100% (v/v) EtOH and Xylene, and embedded in paraffin. Paraffin blocks were cut into 5-µm slices using a HistoCore MULTICUT microtome (Leica Biosystems). Paraffin slices were incubated on a heating plate at 60°C for 30 min, and rehydrated in Xylene (2x 5 min), 100% EtOH (2x 5 min), 96% EtOH (2x 3 min), 70% EtOH (2x 3 min) and water (2x 3 min), followed by antigen retrieval in TRIS-EDTA buffer (pH 9) at 95°C for 30 min. Slices were then washed in 1x PBS with 0.1% (v/v) Tween 20 3x for 5 min and blocked in the buffer with 10% (v/v) goat serum (Sigma-Aldrich) in a humified chamber at RT for one h. Next, slices were incubated with primary antibodies in the buffer in a humified chamber at 4°C overnight. These included mouse anti-E-CAD (1:50, BD Biosciences, 610181), rabbit anti-Villin (1:50, Novus Biologicals, NBP1-85335) and rabbit anti-NTCP (1:50, Novus Biologicals, NBP1-60109). After that, slices were washed in 1x PBS with 0.1% (v/v) Tween 20 3x for 10 min and incubated with secondary antibodies in the buffer, *i.e.* goat anti-mouse-Alexa Fluor 488 (1:100, Thermo Fisher Scientific, A11029) and goat anti-rabbit-Alexa Fluor 647 (1:100, Thermo Fisher Scientific, A21245), and with DAPI (1 μg/mL, Thermo Fisher Scientific) in a humified chamber in the darkness at RT for one h. Then, slices were washed in 1x PBS with 0.1% (v/v) Tween 20 3x for 10 min in the darkness, followed by mounting on Superfrost Plus slides (Thermo Fisher Scientific) with ProLong Diamond Antifade Mountant (Thermo Fisher Scientific) and #1,5 thickness (0,17 mm) cover glass, and immediate imaging. For imaging, an inverted Leica TCS SP8 confocal microscope and the LAS X software (Leica Microsystems) were used. Laser power of 70% was applied. The frame or stack sequential mode from the dye assistant for DAPI, Alexa Fluor 488 and Alexa Fluor 674 was used for excitation and detection without crosstalk between the channels. The detectors used were PMT for DAPI and HyD for Alexa Fluor 488 and Alexa Fluor 647. The objectives used were HC PL FLUOTAR L 20x/0.40 DRY or HC PL APO 40x/0.95 DRY. Images were acquired using the acquisition mode xyz, a pinhole 1 airy unit, format 2048 x 2048 and speed 100. Line average 2-4 for a strong but blurry signal, and line accumulation 2-4 for a weak but sharp signal were applied. An inverted Leica Thunder DMi8 fluorescence microscope and the LAS X software (Leica Microsystems) were used to generate and merge tile scans of entire whole-mount-stained 1.5-cm HFM pieces. Excitation and detection were applied for DAPI and Alexa Fluor 488. The objective used was HC PL FLUOTAR 10x/0.32 DRY. A computational clearing was applied.

### A calcein-AM assay

To examine the confluency of HFM-grown monolayers, the system was incubated with 1 μM calcein-AM (Santa Cruz Biotechnology) in EM in a humidified atmosphere at 37°C and 5% CO_2_ for 15 min. Monolayers were washed 3x in EM and immediately imaged using the FITC channel of an EVOS M5000 Imaging System (Thermo Fisher Scientific). For the functional efflux assay, differentiated monolayers was pre-incubated in DM with 10 μM PSC-833, 10 μM KO-143 and 50 μM MK-571 (inhibitors, Selleck Chemicals) or with an equivalent amount of DMSO in a humidified atmosphere at 37°C and 5% CO_2_ for one h. Then, monolayers were incubated in DM with the inhibitors or DMSO and with 1 μM calcein-AM in a humidified atmosphere at 37°C and 5% CO_2_ for 15 min. Monolayers were washed 3x in DM and immediately imaged using the FITC channel of an EVOS M5000 Imaging System (Thermo Fisher Scientific). The fluorescence was quantified using the ImageJ software (https://imagej.net/ij/).

### Chip assays

A biochamber was printed in-house in the same way as the holder. The design is also the subject of a patent process. The holder carrying HFM-grown monolayers was washed once in HBSS (with calcium, magnesium, glucose and sodium bicarbonate; pH 7-7.4; Thermo Fisher Scientific) supplemented with 1% (v/v) GlutaMAX, 10 mM HEPES and 1 mM sodium pyruvate (Thermo Fisher Scientific), then placed in the biochamber and sealed with rubber O-rings. 710 μl buffer was added to the apical compartment of the biochamber, and sampling ports were sealed with luer locks. For 10-min perfusions (basal flow-through), the biochamber was placed on a heating plate at 37°C and connected to an Ismatec IPC peristaltic pump (Masterflex) using the essential Slang TYGON LMT-55 (PVC; internal diameter 0.51 mm, outer diameter 2.31 mm; Roth) tubing and an additional short tubing (silicone; internal diameter 1.65 mm, outer diameter 3.35 mm; Avantor). For 3-h perfusions (basal circulation), the two latter types of tubing were used together with an additional Teflon tubing (internal diameter 1 mm, outer diameter 3 mm; 2700, Rubber) to minimize unspecific binding of hydrophobic compounds. For 3-h perfusions, a subsampling of 200 or 400 μl circulation buffer was performed and 200 or 400 μl fresh buffer was added. After perfusions, the biochamber was disassembled, and fluids from the apical compartment and basal circulation were collected for the quantification of the compounds using LC-MS/MS or for the quantification of FITC-dextran (FD4, 4000 g/mol, 50 μM; Sigma-Aldrich) or FITC-inulin (3500 g/mol, 0.1 mg/mL; Sigma-Aldrich) fluorescence. For the FD4 fluorescence quantification, 100 μl fluid was added to a 96-well black-walled plate (Agilent) and measured using the FITC channel of a CLARIOstar Plus multi-mode microplate reader (BMG LABTECH). For the FITC-inulin fluorescence quantification, transparent 96-well plates (CELLSTAR) and a GloMax Discover microplate reader (Promega) were used. Intrinsic clearance of midazolam (CL_int_) and apparent permeability P_app_ of FD4 were calculated using standard formulas.^[36,37]^ Small intestine-like organoid monolayers were cultured on Transwell membranes, as described previously^[38]^. Monolayers were exposed on the apical/luminal-like side. Basal effluents were collected after one and two hrs, and combined for subsequent perfusion of the hepatocyte-like system for three hrs. Kidney proximal tubule-like HFM-grown monolayers were established from the immortalized cell line ciPTEC14.4 that displays the active uptake and efflux transport^[17]^. These monolayers were processed for perfusion in the same way as hepatocyte-like monolayers. To further minimize the unspecific binding of hydrophobic compounds during 3-h perfusions, the tubing was first saturated overnight with the buffer containing the parent dose of each compound.

### Liquid chromatography-mass spectrometry/mass spectrometry (LC-MS/MS)

The drug-metabolizing activity of HFM-grown monolayers and ICOs was evaluated using a cocktail of parent compounds, as described previously^[31]^. These included 5 μM midazolam (BUFA; metabolites: 1’-hydroxymidazolam, Sigma-Aldrich; and 1’-hydroxymidazolam glucuronide, LGC Standards), 15 μM dextromethorphan (Santa Cruz Biotechnology; metabolite: dextrorphan tartrate, Sigma-Aldrich), 20 μM tolbutamide (Sigma-Aldrich; metabolite: 4-hydroxytolbutamide, LGC Standards), 15 μM phenacetine (Sigma-Aldrich; metabolite: acetaminophen, Sigma-Aldrich), 20 μM bupropion-HCl (Sigma-Aldrich; metabolite: hydroxybupropion, Sigma-Aldrich) and 12 μM 7-hydroxycoumarin (umbelliferone, Sigma-Aldrich; metabolite: 7-hydroxycoumarin glucuronide, LGC Standards). Differentiated monolayers and organoids (7 days) were exposed with the cocktail in DM in a humidified atmosphere at 37°C and 5% CO_2_ for 24 hrs. Collagen I *sandwich* PHHs were previously cultured for 48 hrs and then exposed for four hrs,^[31]^ and data were extracted and normalized. For organoids and PHHs, 1’-hydroxymidazolam glucuronide was not measured. For liver-on- a-chip experiments, 5 μM midazolam in the buffer was used. For double chip experiments, hepatocyte-like and Transwell monolayers were exposed to 50 μM midazolam, 50 μM coumarin (Sigma-Aldrich) and 50 μM metformin (Sigma-Aldrich) in the buffer. For a CYP3A4 inhibition assay, HFM-grown monolayers were incubated in DM with 10 μM ketoconazole (Selleck Chemicals) or an equivalent amount of DMSO for three hrs and then in DM with 1 μM ketoconazole or DMSO and 5 μM midazolam for 24 hrs. 200 μl of culture medium or buffer were collected in the 1.5-mL Short Thread Vials with the ND9 Short Thread Screw Caps (BGB Analytik) containing 200 μl ultrapure MeOH, vortexed and stored at -20°C until the quantification of the parent compounds and metabolites using LC-MS/MS. As described previously,^[31]^ samples were centrifuged for 10 min at 1500 g to precipitate any protein prior to the analysis. Standards of the parent compounds and metabolites were prepared in the same matrix (medium or buffer) as samples. Usually, 1 μl medium or buffer with MeOH was injected. If necessary, selected metabolites were quantified using a high sensitivity method with a 5-μl injection. Standards and samples were analyzed in a single run using a Shimadzu triple-quadrupole LCMS 8050 system with two Nexera XR LC-20AD pumps, a Nexera XR SIL-20AC autosampler, a CTO-20AC column oven and an FCV-20AH2 valve unit (Shimadzu). The compounds were separated on a Synergi Polar-RP column (150 × 2.0 mm, 4 µm, 80 Å) with a 4 × 2 mm C18 guard column (4 × 2 mm, Phenomenex). The mobile phase consisted of 0.1% (v*/*v) formic acid in ultrapure water (A) and 0.1% (v*/*v) formic acid in MeOH (B), and was set as 100% A (0-1 min), 100% to 5% A (1-8 min), 5% A (8-9 min), 5% to 100% A (9-9.5 min) and 100% A (9.5-12.5 min). The total run time was 12.5 min, and the flow rate was 0.2 mL/min. Peaks were integrated using the LabSolutions software (Shimadzu). For each compound, a limit of quantification (LOQ) was determined based on a standard curve and a limit of detection (LOD) was determined based on a noise signal. For each sample set, a dose (time 0 hrs) and a blank (negative control) were measured. For each parent compound and its metabolite(s), data were normalized to an actual (measured) dose in the relation to an expected (calculated) dose. Intracellular levels of the parent compounds and metabolites were not taken into account. Efflux of metformin was normalized to total FD4 fluorescence in the apical compartment (permeability).

### Statistical analysis

All descriptive statistical analyses were performed using the Microsoft Excel (Microsoft) and GraphPad Prism software (GraphPad), unless otherwise stated. All data are presented as mean ± SD (error bars) with individual independent (donors) or technical replicates marked, unless otherwise stated. Descriptive statistical details of the experiments can be found in the figure legends. For experiments with three or four independent replicates, the non-parametric Kruskal-Wallis test followed by the Dunn’s multiple comparisons’ test were performed. Even if medians varied significantly in the Kruskal-Wallis test (P-value < 0.05), no statistically significant differences were found in the Dunn’s test (adjusted P-value < 0.05) due to high inter-donor variance (SD). Thus, trends are presented.

## Results

### Organoid fragments form polarized hepatocyte-like monolayers on laminin 332- coated HFMs

To enable the formation of cell monolayers, HFMs were coated with biological polymeric glue, L-DOPA, that facilitates subsequent coating with ECM components^[14,17–19]^. To replace animal-derived and undefined ECM components such as Matrigel, we examined recombinant human laminins that represent the entire spectrum of α1-5 chains crucial for binding to respective integrins, *i.e.* laminin 111, 211, 332, 411 and 511^[39]^. Coated HFMs were seeded with fragments of organoids that were grown in expansion medium in standard Matrigel droplet-based cultures (Fig. S1A) and can be matured into hepatocyte-like cells upon a switch to differentiation medium^[21]^ (Fig. S1B). Only organoid fragments grown on laminin 332- and 511-coated HFMs succeeded in generating confluent monolayers over 7 days (Fig. S2A). Upon differentiation, the expression of crucial functional markers of mature hepatocytes, *i.e. ALB* (albumin), phase I CYP450 (*CYP3A4*, *CYP2C9*, *CYP2D6*, *CYP2B6*) and phase II glucuronosyltransferase/UGT and sulfotransferase/SULT (*UGT2A3*, *SULT1A1*) drug-metabolizing enzymes as well as apical *ABCB1* (MDR1) and basolateral *ABCC3* (MRP3) drug transporters was upregulated in monolayers grown on laminin 332-coated HFMs, as compared to laminin 511-coated HFM-grown monolayers and Matrigel organoid cultures from matched donors (Fig. S2B). However, the expression for *ALB*, *CYP2D6* and *CYP2B6* was downregulated in laminin 332-coated HFM-grown monolayers versus the whole adult liver from matched donors. As laminin 332 drives both efficient expansion and maturation, it was selected for further developments (Fig. 1A).

**Fig. 1.**
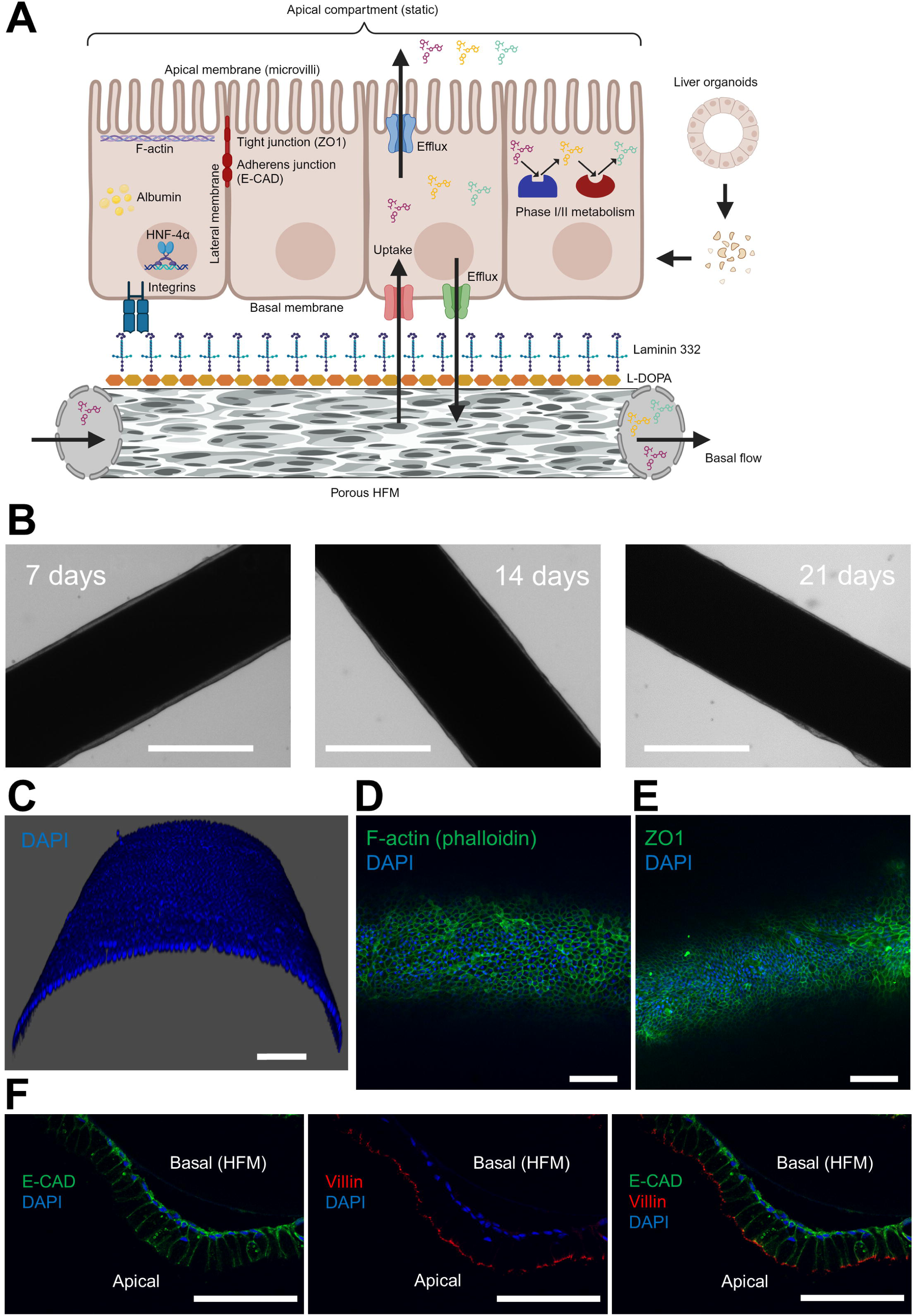
Establishing hepatocyte-like HFM-grown monolayers optimized for organ-on-a-chip assays. (A) A scheme showing a minimal viable product for the liver-on-a-chip system. (B) Representative brightfield images of confluent monolayers (a median section) after 3 days in pre-DM, and 7, 14 and 21 days in DM under the static culture. (C) A representative 3D-reconstituted whole-mount confocal immunofluorescence microscopy image of a HFM-grown monolayer (an anterior-dorsal section) after 7 days in DM. (D) A representative whole-mount confocal immunofluorescence microscopy image of F-actin (phalloidin) in a dorsal section of a HFM. (E) A representative whole-mount confocal immunofluorescence microscopy image of ZO-1 in a dorsal section of a HFM. (F) A representative confocal immunofluorescence microscopy image for the basolateral localization of E-CAD and the apical localization of Villin in cell membranes of a monolayer in an anterior section of a HFM. Data information: scale bars; in B, 500 µm; in C-F, 100 µm. In A, the scheme was prepared using Biorender (https://www.biorender.com/). In C-F, nuclei were counterstained with DAPI. In B, three independent replicates (donors; 2, 3, 5) with six technical replicates (monolayers) were examined; in C-F, three independent replicates (donors; 2, 3, 5) with three technical replicates (monolayers) were examined.

We did not observe any morphological changes when differentiated monolayers were monitored for 7, 14 and 21 days (Fig. 1B), demonstrating the potential for long-term studies. A 3D reconstruction revealed that monolayers were intact (Fig. 1C). In addition, we detected a ubiquitious presence of cytoskeleton-associated F-actin (Fig. 1D and S3) and tight junction-associated ZO1 (Fig. 1E) in cell membranes. To enable passive diffusion (transcellular and paracellular) and active transport of drugs^[12]^ between two microfluidic compartments, monolayers must form columnar tubular epithelium with apical-to-basolateral polarization, as opposite to the native hepatic canalicular architecture with the apical compartment between cells^[40]^. We confirmed the basolateral localization of adherens junction-associated E-CAD and the apical localization of microvilli-associated Villin (Fig. 1F), and the (sub)apical localization of F-actin (Fig. S4). Another sign of cell polarization was a clear localization of nuclei along basal membranes (Fig. 1F). Thus, monolayers exhibited inside-outside polarization, *i.e.* the apical side towards outside and the basal side towards inside the tube (lumen). Together, laminin 332 mediated expansion, hepatocyte-like differentiation and polarization of organoid cell monolayers on HFMs.

### Hepatocyte-like HFM-grown monolayers display a higher expression of drug metabolism- and transport-relevant genes, as compared to standard organoid cultures

We compared the total transcriptome of HFM-expanded monolayers and HFM-expanded monolayers differentiated for 7, 14 and 21 days with both expanded and differentiated standard Matrigel droplet-based organoid cultures, and with the whole liver, all from matched or partly matched donors. A principal component analysis revealed that monolayer and organoid transcriptomes are substantially separated (about 8.5 units) from the liver sample cluster along the first principal component (PC1) that describes the most variation in data (Fig. 2A). Based on the second principal component (PC2) that illustrates the second largest variation of transcriptomes, we identified clear clusters of expanded and differentiated monolayers and organoids. mRNA profiles of differentiated organoids and expanded monolayers were similar and separated from differentiated monolayers by up to three units. The most distant cluster from differentiated monolayers were expanded organoids. We did not detect distinct clusters for the three different time points of differentiated monolayers, indicating gene expression stability over time. To further depict transcriptomic differences between differentiated monolayers and organoids, we conducted a functional annotation analysis using the bioinformatic tool *DAVID*. Genes upregulated in monolayers over organoids encode proteins that are involved in numerous biological processes active in mature hepatocytes, including retinol metabolism, drug/xenobiotic metabolism, lipid transport and metabolism, albumin secretion, aldehyde dehydrogenase activity and gluconeogenesis (Fig. 2B).

**Fig. 2.**
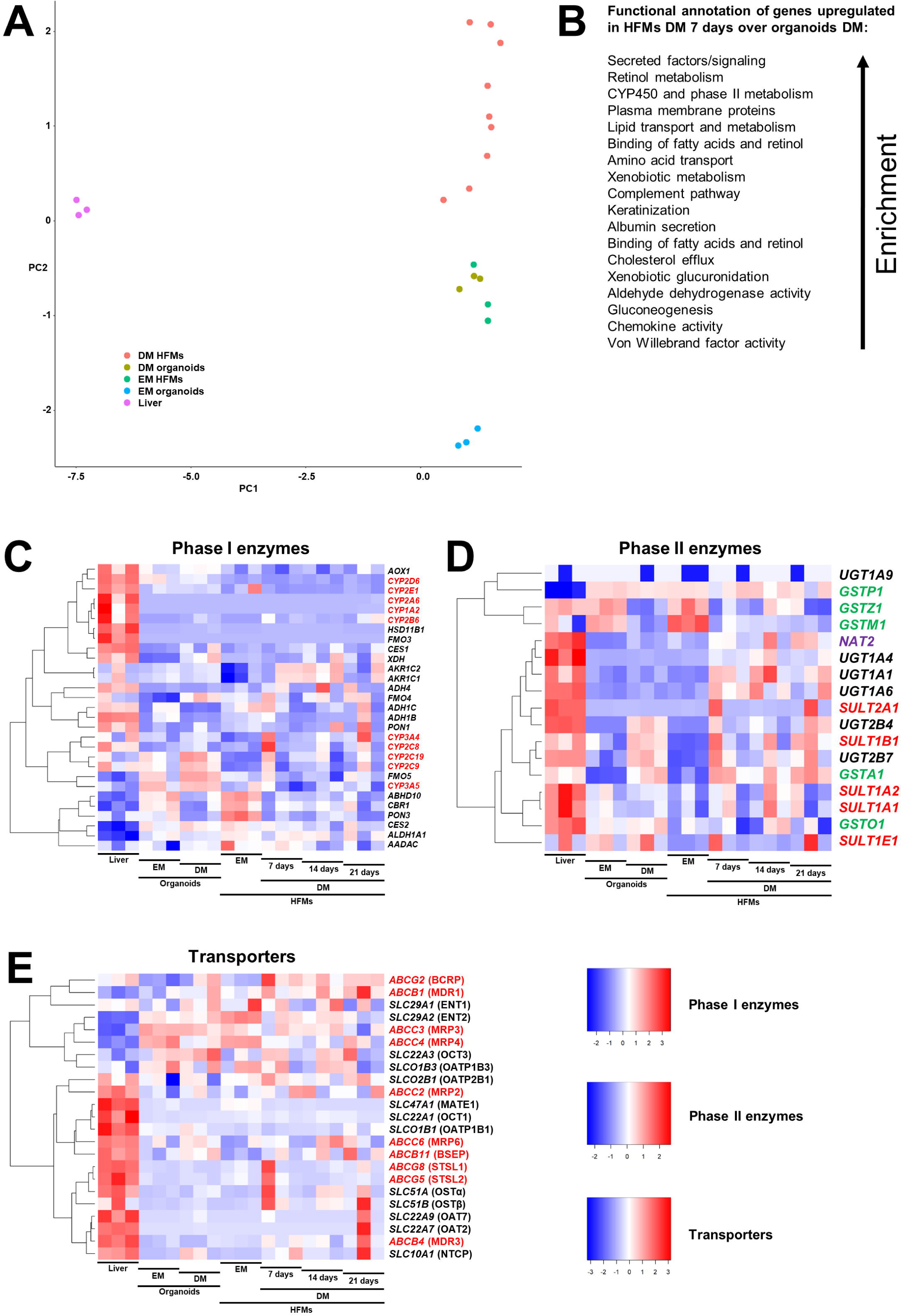
Bulk mRNA sequencing of PK/ADME-relevant genes in statically cultured HFM-grown monolayers, as compared to organoids and liver tissue. (A) A principal component analysis. (B) A functional annotation of genes upregulated in monolayers over organoids. (C) A heatmap for gene expression of phase I drug- metabolizing enzymes. (D) A heatmap for gene expression of phase II drug-metabolizing enzymes. (E) A heatmap for gene expression of drug transporters. Data information: in C, CYP450 and non-CYP450 enzymes are marked in red and black, respectively. In D and E, different enzyme or transporter families are marked with different colors. In C-E, the heatmaps show normalized log2 gene expression. A 5% FDR (adjusted P-value < 0.05) cut-off was applied. Values were scaled (z-score by row) with breaks from ≤ -2 to ≥ 3 in C, ≤ -2 to ≥ 2 in D and ≤ -3 to ≥ 3 in E. In A-E, three independent replicates per condition were examined. In A-E, matched donors (2, 3, 5) were used for organoids and monolayers. In A and C-E, donor 3 matched liver tissue, organoids and monolayers, and two other donors (1, 4) were used for liver tissue. The order of the donors: 3, 1, 4 for liver tissue; 2, 3, 5 for organoids (EM and DM); 2, 5, 3 for monolayers EM and DM 14 days; 3, 5, 2 for monolayers DM 7 days; 5, 3, 2 for monolayers DM 21 days.

Next, we focused on the expression of individual genes encoding proteins that participate in metabolism and transport of drugs. First, we examined the abundance of transcripts of phase I drug-metabolizing enzymes, both of the CYP450 (oxidases)^[41]^ and non-CYP450 (oxidases, reductases, hydrolases) family^[42]^ (Fig. 2C). The first cluster encompassed enzymes that were downregulated in differentiated monolayers (7, 14 and 21 days) over the liver, *CYP2D6*, *CYP1A2*, *CYP2B6* and *CES1*, for instance. The second cluster included enzymes that were partly expressed in differentiated monolayers at similar levels to the liver such as *CYP3A4*, *CYP2C9*, *ADH4* and *PON1*. Of note, *CYP3A4*, the most important phase I CYP450 enzyme for drug metabolism,^[41]^ was upregulated in differentiated monolayers over differentiated organoids. The last cluster encompassed non-CYP450 enzymes that were upregulated in monolayers and/or organoids (both expanded and differentiated) versus the liver, including *FMO5*, *ABHD10*, *CBR1* and *ALDH1A1*. Furthermore, we analyzed gene expression of phase II drug-metabolizing enzymes that drive conjugation reactions, *i.e.* UGTs (glucuronidation),^[43]^ SULTs (sulfation),^[44]^ NAT2 (acetylation)^[42]^ and GSTs (glutathione conjugation)^[45]^ (Fig. 2D), which was similar to the liver in differentiated monolayers of some donors. In addition, *UGT1A1*, *UGT1A6*, *SULT1A1* and *NAT2* were upregulated in differentiated monolayers over differentiated organoids. Finally, gene expression analysis of drug transporters^[40]^ revealed a cluster that was upregulated in monolayers and organoids (both expanded and differentiated) over the liver, including basolateral uptake transporters such as *SLCO1B3* (OATP1B3, anionic drugs) and *SLC22A3* (OCT3, cationic drugs), basolateral efflux transporters, *ABCC3* (MRP3) and *ABCC4* (MRP4), and apical efflux transporters, *ABCG2* (BCRP) and *ABCB1* (MDR1) (Fig. 2E). The expression of *SLCO2B1* (OATP2B1, basolateral uptake of anionic drugs) in differentiated monolayers was similar to the liver. *SLC10A1* (NTCP, basolateral uptake), *ABCC2* (MRP2, apical efflux) and *ABCB4* (MDR3, apical efflux) were expressed in some differentiated monolayers at the levels observed in the liver. Other transporters such as *SLCO1B3* (OATP1B1, basolateral uptake of anionic drugs), *SLC22A1* (OCT1, basolateral uptake of cationic drugs) and *SLC47A1* (MATE1, apical efflux) were downregulated in differentiated monolayers versus the liver. In line with the principal component analysis, long- term differentiation of monolayers did not clearly alter gene expression of drug-metabolizing enzymes and transporters. Together, gene expression in differentiated monolayers was more comparable to liver tissue, as compared to standard differentiated organoids from matched donors.

### Hepatocyte-like HFM-grown monolayers exhibit functional transport and metabolism of drugs

We observed a ubiquitious or an abundant membranous presence of representative transporters for basal uptake (OATP1B3 and OCT1) or efflux (MRP3) and for apical efflux (MDR1 and MRP2) in differentiated (7 days) HFM-grown monolayers (Fig. 3A). In addition, the basolateral localization of ubiquitiously expressed NTCP (uptake) was confirmed (Fig. 3B), in line with the previous data on monolayer polarization (Fig. 1F). It was possible to detect a high protein expression of OCT1, despite lower transcript levels, as compared to the liver. For the functional examination of selected transporters, we used calcein-AM that enters cells via passive diffusion and is effluxed by MDR1 or is first hydrolized intracellularly into the fluorescent anion calcein that is effluxed by BCRP and MRPs^[46,47]^. Blocking efflux of calcein-AM and calcein with a cocktail of reference inhibitors, *i.e.* PSC-833 (MDR1), KO143 (BCRP) and MK-571 (MRPs), resulted in a 5.3-fold increase in intracellular fluorescence of calcein (Fig. 3C and D), confirming that the efflux transporters were active.

**Fig. 3.**
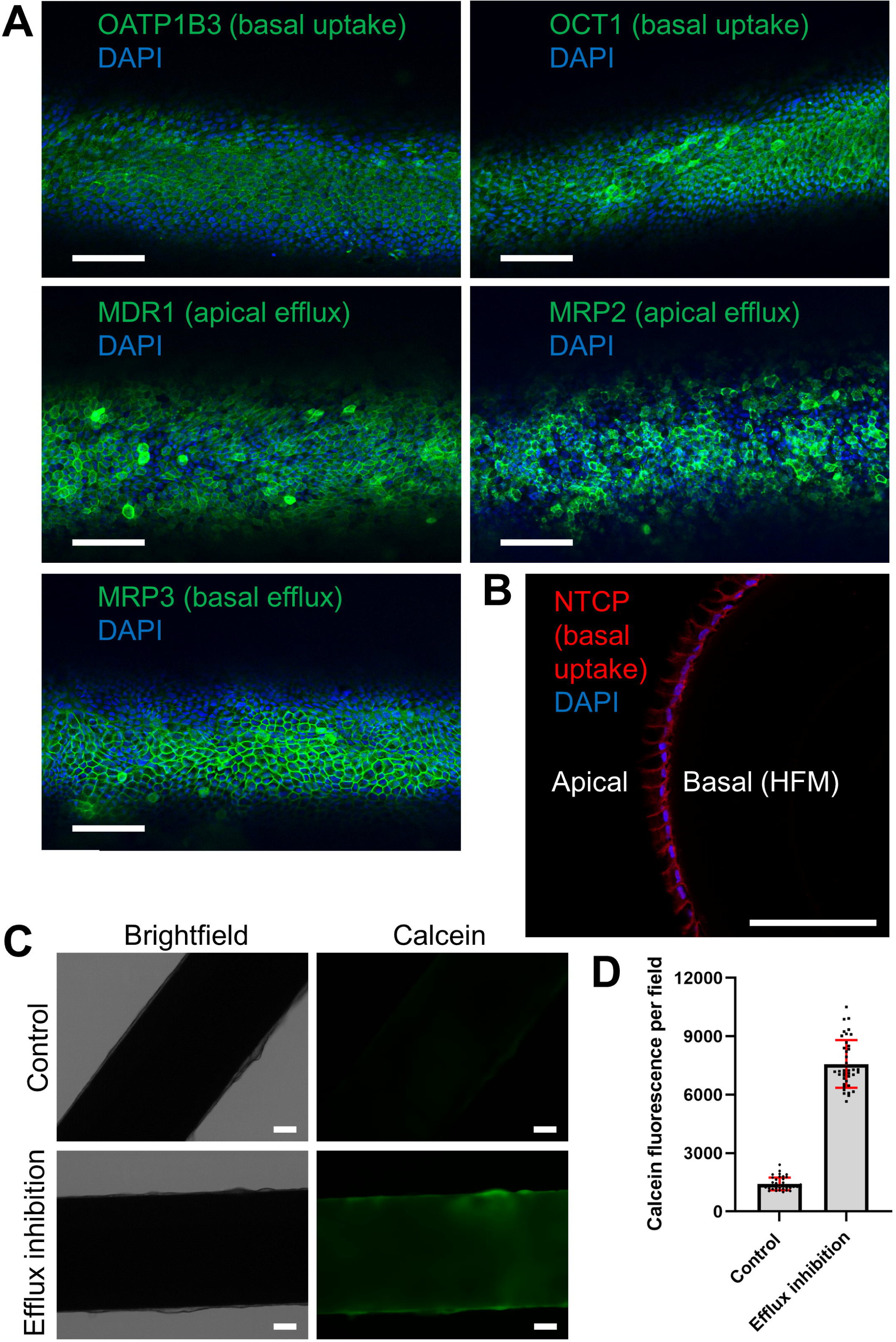
The expression and functionality of drug transporters in cell membranes of statically cultured HFM-grown monolayers. (A) A representative whole-mount confocal immunofluorescence microscopy image of OATP1B3, OCT1, MDR1, MRP2 and MRP3 in a dorsal section of a HFM. (B) A representative confocal immunofluorescence microscopy image for the basolateral localization of NTCP in cell membranes of a monolayer in an anterior section of a HFM. (C) Representative brightfield images and images of intracellular fluorescence of calcein of monolayers treated with efflux inhibitors versus DMSO (control). (D) The quantification of calcein fluorescence. Data information: scale bars in A-C, 100 µm. In A and B, nuclei were counterstained with DAPI. In D, the graph shows mean ± SD (error bars). In A and B, three independent replicates (donors; 2, 3, 5) with three technical replicates (monolayers) were examined. In C and D, one independent replicate (donor, 5) with 40 technical replicates (fields) from two monolayers (20 replicates each) was examined.

To study the oxidative capacity of phase I drug-metabolizing enzymes in differentiated (7 days) HFM-grown monolayers, we examined a cocktail of FDA US reference parent drugs^[48]^ (Fig. 4A). Together, these enzymes mediate approximately 80% of CYP450-associated metabolism of the top 200 most prescribed drugs (in the USA in 2014), with the most important CYP3A4 being responsible for 40%.^[41]^ To examine phase II metabolism via glucuronidation by various UGTs, mainly UGT1A6,^[49]^ 7-hydroxycoumarin (coumarin-OH) was used, which is the major phase I metabolite (via CYP2A6) of coumarin, a natural product (nutraceutical)^[50]^. Glucuronidation is the major phase II reaction, and UGTs jointly carry out 45% of non-CYP450-associated metabolism of the top 200 most prescribed drugs.^[41]^ Metabolism-driven clearance of midazolam and coumarin in the liver, via the so-called first- pass effect, is rapid and extensive. Clearance of dextromethorphan and phenacetin is moderate. In turn, tolbutamide and bupropion are considered low clearance drugs.^[50–52]^ We compared the metabolizing performance of monolayers with standard organoid cultures from matched donors and PHH *sandwich* collagen I cultures (Fig. 4B). We detected metabolites for all the parent compounds in all monolayer cultures, in contrast to organoid cultures with a limited metabolizing capacity. The most extreme case was the lack of bupropion-OH formed in all organoid cultures. The production of midazolam-OH in monolayers was at a similar level to PHH cultures. However, PHHs outperformed monolayers in the formation of the remaining metabolites.

**Fig. 4.**
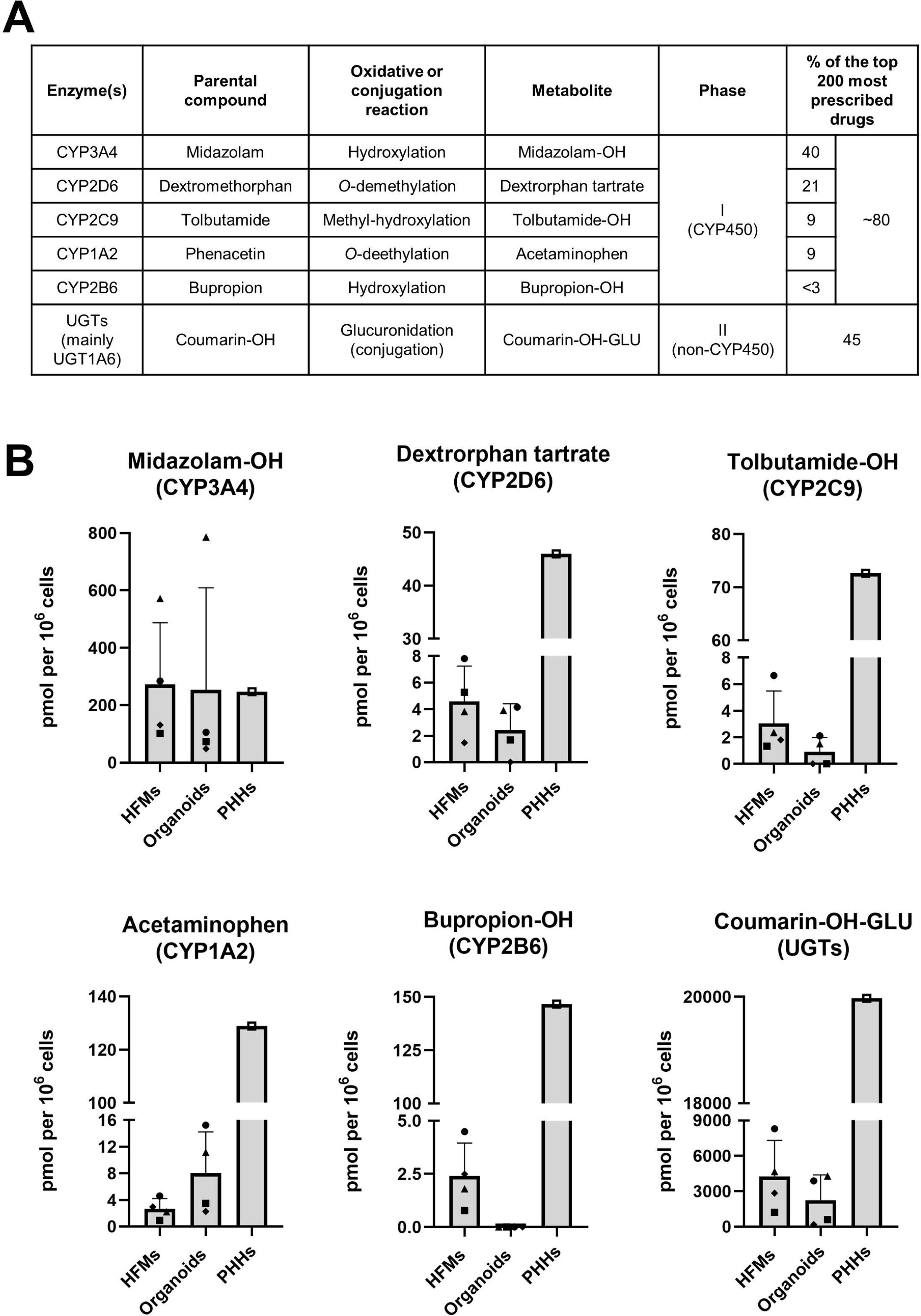
Phase I and II drug metabolism in statically cultured HFM-grown monolayers. (A) Characteristics of selected parent compounds used for testing the activity of the most important phase I and II drug-metabolizing enzymes. (B) The formation of metabolites of the parent compounds in monolayers, as compared to organoids and collagen I *sandwich* PHHs. Data information: in B, the graphs show mean + SD (error bars). Four independent matched replicates (donors; 1-3, 5) were examined for monolayers and organoids. For monolayers, one technical replicate (well/measurement, from pooled six monolayers); and for organoids, two averaged technical replicates (wells/measurements) were examined per donor. PHHs are from a pool of 10 independent replicates (donors, five males and five females) with three technical replicates (wells/measurements).

Next, we explored the biotransformation kinetics of midazolam over 24 hrs in donor 3 monolayers with the highest conversion and donor 5 monolayers with the lowest metabolizing capacity (Fig. 4B). The major primary metabolite of midazolam, 1’- hydroxymidazolam (midazolam-OH), is further transformed into 1’-hydroxymidazolam glucuronide (midazolam-OH-GLU) by two other UGT enzymes, *i.e.* UGT2B4 and UGT2B7.^[53]^ Donor 3 monolayers showed a 7-fold higher capacity to form midazolam-OH and a 14-fold increase in the formation of midazolam-OH-GLU after 24 hrs, as compared to donor 5 monolayers (Fig. S5A). In addition, the production of both midazolam-OH and midazolam- OH-GLU in donor 5 monolayers reached saturation already after 16 hrs. These findings were reflected by an upregulated expression of *CYP3A4*, *UGT2B4* and *UGT2B7* in donor 3 over donor 5 monolayers (Fig. 2C and D).

The interaction between midazolam and ketoconazole is a classical example of a drug-drug interaction. Ketoconazole can compete for CYP3A4, compromising the conversion of midazolam to midazolam-OH.^[54]^ Upon exposure to ketoconazole, we demonstrated a reduced formation of midazolam-OH and midazolam-OH-GLU in both donor 3 and 5 monolayers (Fig. S5B). Together, the hepatocyte-like HFM-based system allows for studying phase I and II metabolism of drugs in a personalized fashion by capturing donor-to-donor variations in the activity of biotransforming enzymes. Moreover, the system is suitable to examine active transport of drugs across cell membranes.

### Hepatocyte-like HFM-grown monolayers are suitable to study transport and metabolism of drugs on-a-chip

The ultimate step in advancing our model was coupling it to a micro-/millifluidic system using biocompatible in-house-3D printed chips. We grew and differentiated (7 days) organoid fragments on HFMs immobilized on inserts that were then placed into biochambers. By this, two microfluidic compartments were created, *i.e.* a perfusable basal/blood-like compartment inside HFMs and a static apical/canalicular-like compartment outside HFMs. We perfused the system with FITC-dextran (FD4) solution for 10 min with a speed of 50 µl/min (3 mL/h) (Fig. 5A) that is within the standard range of 2-6 mL/h applied for HFM-based systems previously^[14,17–19]^. We confirmed the functional barrier integrity of monolayers of both donors, as demonstrated by 10 times lower paracellular passive diffusion of FD4 to the apical compartment, *i.e.* apparent permeability (P_app_)^[37]^, as compared to cell-free HFMs (coated with L-DOPA and laminin 332, a positive control) (Fig. 5B). Nail polish-coated HFMs served as a tight barrier (a negative control).

**Fig. 5.**
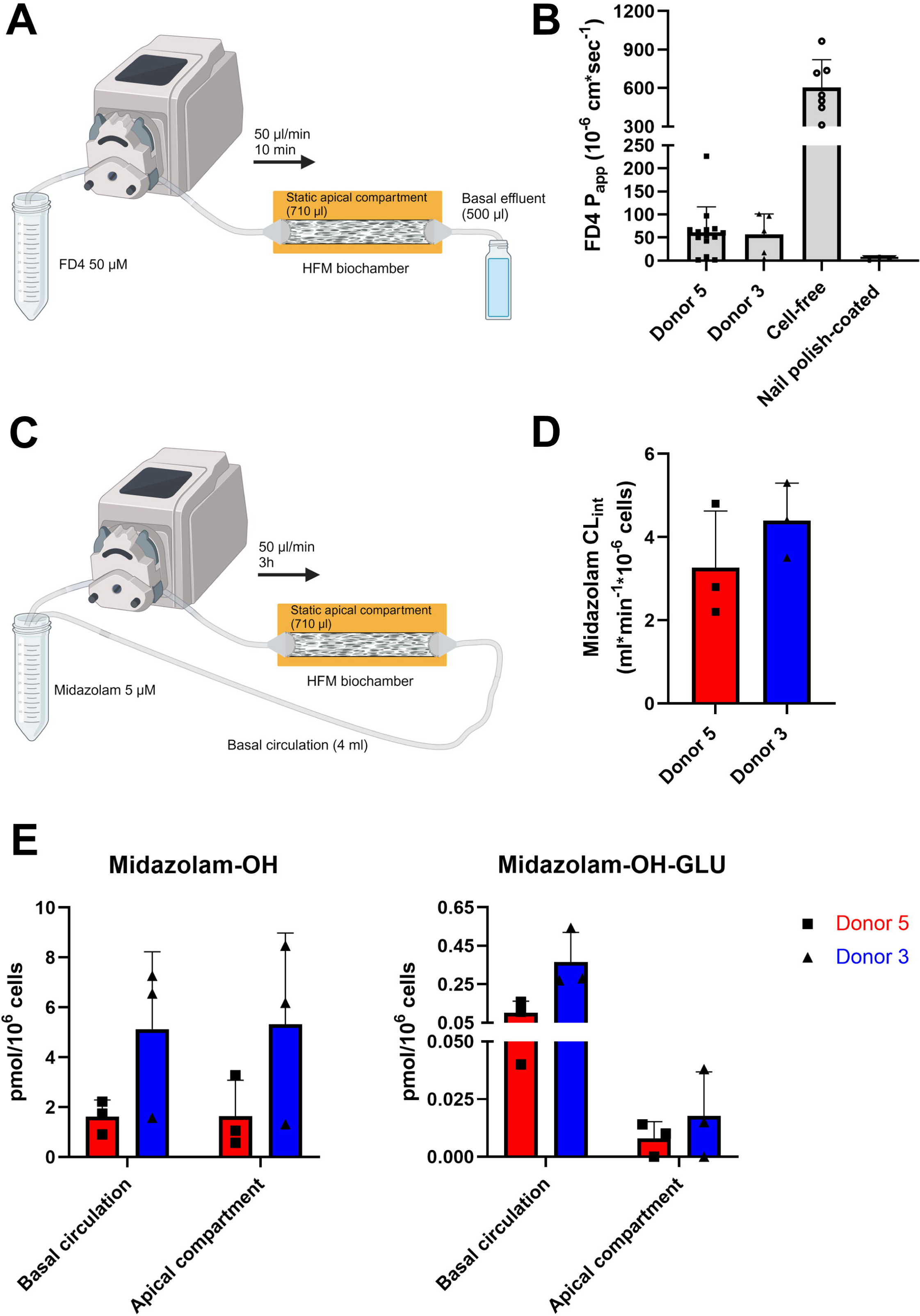
The functional barrier integrity of HFM-grown monolayers and the disposition of midazolam on-a- chip. (A) A scheme for the 10-min perfusion setup. (B) FD4 P_app_ of monolayers, as compared to cell-free HFMs (a positive control) and nail polish-coated HFMs (a negative control) after 10-min perfusion. (C). A scheme for the 3- h perfusion setup. (D) CL_int_ of midazolam in monolayers over 3-h perfusion. (E) The disposition of midazolam-OH and midazolam-OH-GLU in the basal circulation and static apical compartment after 3-h perfusion. Data information: in A and C, the schemes were prepared using Biorender (https://www.biorender.com/). In B, D and E, the graphs show mean + SD (error bars). In E, midazolam-OH concentrations were ≥ LOQ and midazolam-OH- GLU concentrations were ≤ LOQ and ≥ LOD. In A and B, two independent replicates (donors; 3, 5) with five and 14 technical replicates (chips) were examined, respectively. Seven and three independent replicates of cell-free and nail polish-coated HFMs were analyzed, respectively. In C-E, two independent replicates (donors; 3, 5) with three technical replicates (chips) were examined.

Next, the disposition of midazolam was studied on-a-chip. To minimize the so-called dead volume and thus to enable detection of the metabolites,^[10]^ we designed a circulation system with a 4-mL volume (Fig. 5C). Over three hours, donor 3 monolayers metabolized 42% and donor 5 monolayers 37% of midazolam (Fig. S6A). Based on linear fragments of logarithmic depletion curves of midazolam (Fig. S6B), we determined slopes that were necessary to calculate its intrinsic clearance (CL_int_)^[10,36]^ (Fig. 5D). CL_int_ for donor 3 monolayers was 1.3- fold higher, as compared to donor 5 monolayers. In line with the data generated on the statically cultured system, donor 3 monolayers produced higher amounts of midazolam-OH and midazolam-OH-GLU, as compared to donor 5 monolayers (Fig. 5E). For both donor monolayers, we observed an equilibrium of midazolam-OH between the basal and apical compartment. In turn, we detected higher amounts of midazolam-OH-GLU in the basal compartment for both donor monolayers, likely explained by the presence of MRP3 (Fig. 3A) pumping the hydrophilic metabolite out to the circulation^[55]^ for renal excretion^[53]^. A morphological evaluation of monolayers after perfusion did not reveal any visible damage (Fig. S7).

Finally, we established a proof-of-concept for a multi-organ-on-a-chip platform by perfusing the hepatocyte-like system for three hrs with a basal effluent from an organoid-based small intestine-like static Transwell system^[38]^ to study the first-pass effect and bioavailability of midazolam and coumarin upon oral administration (apical/luminal-like compartment) (Fig. 6A), similar to previous double-chip studies using different compounds^[56–59]^. The most efficient metabolizer was used, donor 3 monolayers. Midazolam-OH was produced by both small intestine- and hepatocyte-like monolayers (Fig. 6B), in line with CYP3A4 expression in the small intestine^[38]^. However, we did not detect midazolam-OH-GLU due to a low amount of midazolam that diffused to the basal effluent of the small intestine-like system, *i.e.* 3.4%. Ultimately, we found 2.3% of the midazolam dose in the basal compartment of the hepatocyte-like system (bioavailability). In contrast, coumarin was metabolized to coumarin- OH, followed by its glucuronidation, in the hepatocyte-like system only thanks to an exclusive activity of the minor phase I enzyme CYP2A6^[41,48]^ (Fig. 6C). Detection of coumarin-OH was possible despite a lower expression of *CYP2A6* in comparison to PHHs (Fig. 2C). The bioavailability of coumarin was 3.3%. Moreover, we connected the hepatocyte-like system to a previously established kidney proximal tubule-like HFM-based system^[17]^ to examine the disposition of the hydrophillic cationic drug metformin that is not metabolized in the liver, but is subjected to renal excretion via active transport to the apical/urine compartment^[46,60]^ (Fig. 6D). After 10-min perfusion, we measured metformin efflux to the apical compartment of both the systems and we found 2.6-fold increased efflux to the kidney proximal tubule-like apical compartment (Fig. 6E). Together, the hepatocyte-like HFM-based system on-a-chip maintains a tight barrier over 3-h perfusion and allows for studying the disposition of midazolam and coumarin.

**Fig. 6.**
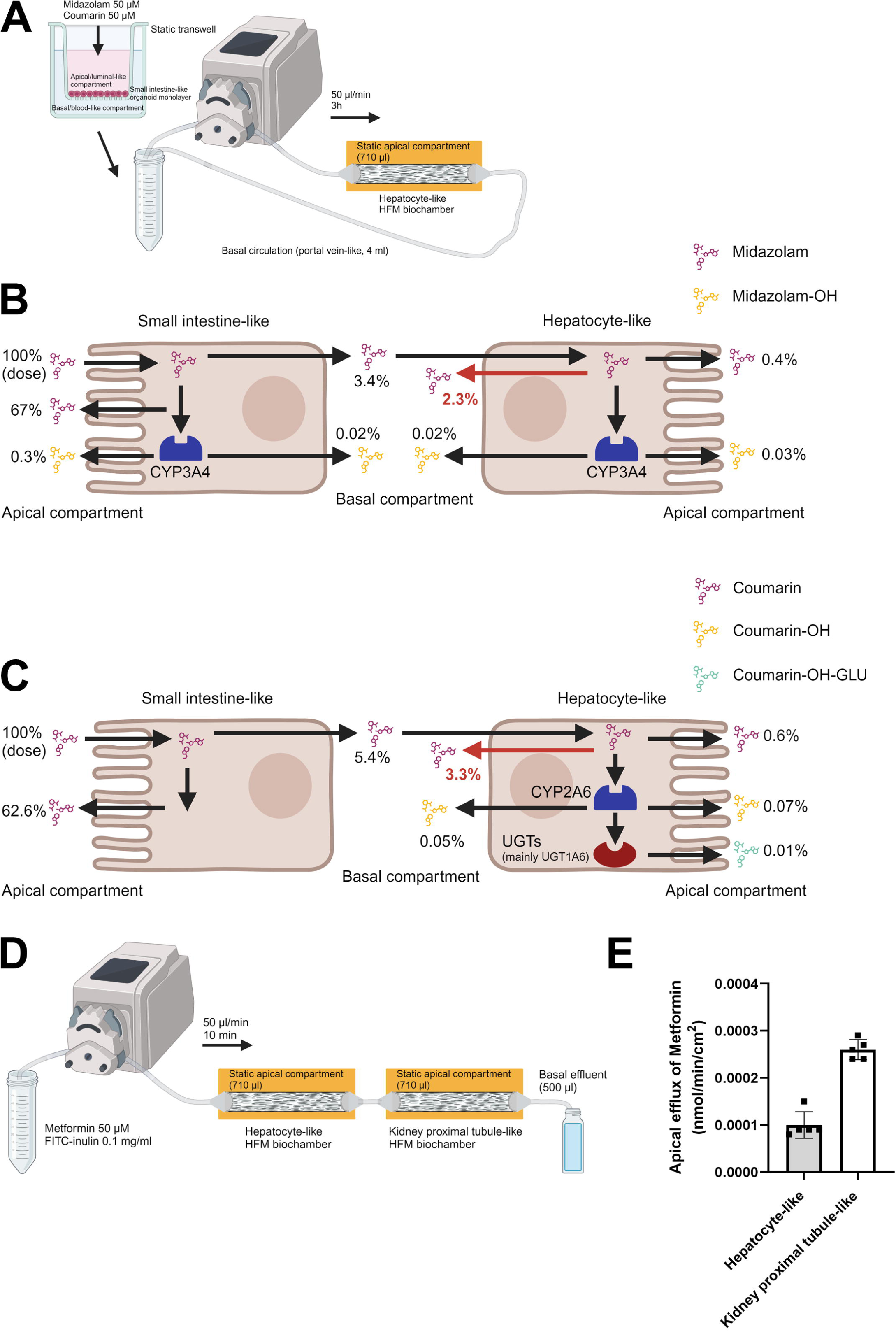
Connecting the liver-on-a-chip system to other PK/ADME-relevant organ systems. (A) A scheme for the connection to the statically cultured small intestine-like Transwell system. (B) The disposition of midazolam and midazolam-OH in the apical compartment and basal effluent of the small intestine-like system, and in the basal circulation and static apical compartment of the hepatocyte-like system after the administration of 50 µM midazolam over two hrs, followed by 3-h perfusion. (C) The disposition of coumarin, coumarin-OH and coumarin- OH-GLU in the apical compartment and basal effluent of the small intestine-like system, and in the basal circulation and static apical compartment of the hepatocyte-like system after the administration of 50 µM coumarin over two hrs, followed by 3-h perfusion. (D) A scheme for the connection to the kidney proximal tubule-like HFM- based system on-a-chip. (E) Efflux of 50 µM metformin to the static apical compartment of the hepatocyte- and kidney proximal tubule-like systems. Data information: In A-D, the schemes were prepared using Biorender (https://www.biorender.com/). In E, the graph shows mean ± SD (error bars). In A-C, one independent replicate (donor, 3) with three technical replicates (chips) was examined; in D and E, one independent replicate (donor, 5; and one passage of the ciPTEC14.4 cell line) with five technical replicates (chips) was examined.

## Discussion

Despite the recent progress in developing human microphysiological systems, robust and animal component-free liver-on-a-chip models that are suitable for examining both metabolism and transepithelial transport of drugs remain an unmet need for preclinical PK/ADME prediction. Here, we filled this gap with the millifluidic hepatocyte-like system based on organoid-derived polarized monolayers grown on HFMs coated with recombinant human laminin 332. Hence, our model allows for studying uptake of parent compounds from the perfusable basal/blood-like compartment, and excretion of those compounds and their metabolites to the static apical/canalicular-like compartment or back to the circulation. In addition, the system displays improved hepatocyte-like maturation and functionality over standard organoid cultures from matched donors, similar to our previous bile duct-on-a-chip HFM-based model^[19]^. We believe that this is the result of a concerted action of the 3D HFM scaffold, organoid cells, liver-specific laminin 332 and growth factors in differentiation medium. Chip systems utilizing human organoid-derived cells are considered one of the most powerful *in vitro* models for preclinical drug development in the future, once standardized and scaled up.^[61]^ Our model pioneers the field of liver-on-a-chip systems both by utilizing organoids and complying to the 3Rs’ principles^[3]^.

We identified laminin 332 as the most efficient driver of expansion of organoid fragments on HFMs, resulting in full coverage, as well as of hepatocyte-like differentiation. Enhancing the proliferative capacity is in line with studies in mice with acute liver injury that showed a dense deposition of laminins critical for hepatocyte progenitor cell-mediated regeneration^[23,62]^. In agreement with it, adult stem cell-derived organoids are believed to recapitulate epithelial regenerative processes during their expansion phase^[63]^. Though, laminin 332 seems the least obvious choice from all commercially available recombinant human laminins for increasing hepatocyte-like maturation, as only one study linked it with the adult liver during homeostatic maintenance^[22]^. Furthermore, this laminin can support cholangiocyte-like differentiation *in vitro*.^[64]^

By using organoids derived from four donors of both sexes, we were able to capture inter-individual variabilities in the expression and activity of drug-metabolizing enzymes and drug transporters. To ensure a personalized approach, generating data from more donors of both sexes and various ages is necessary. An added value will be the knowledge on clinically-relevant genetic polymorphisms of drug-metabolizing enzymes^[65]^ and drug transporters^[40]^ of those individuals. Our liver-on-a-chip system allowed for determining experimental kinetic parameters of drugs and their metabolites upon at least 3-h perfusion. We examined the basal and apical disposition of midazolam that is the substrate for the most important drug-metabolizing enzyme, CYP3A4^[41]^, whose role will even increase in a long-term perspective by driving metabolism of 64% of all newly approved drugs.^[41]^ 3-h perfusion is a progress in comparison to our previous HFM-based systems with perfusion applied of only 10-13 min^[14,17–19]^. To evaluate preclinical predictiveness of our system for PK/ADME, *in vitro* data have to be quantitatively extrapolated into human profiles using *in silico* physiologically-based pharmacokinetic (PBPK) mechanistic modeling^[10,36,66]^. Examining phase II metabolism-related conjugation reactions other than the most frequently detected glucuronidation such as sulfation and acetylation^[52]^, will also be an important element for further development of the model.

Despite our advancements, the drug-metabolizing capacity of the statically cultured system was lower for five out of the six compounds tested, as compared to PHHs, with only the formation of midazolam-OH being at the similar level. However, the liver-on-a-chip system exhibited three or six times reduced clearance of midazolam, as compared to PHHs in two existing millifluidic circulation systems^[10,36]^. To overcome these limitations, cell differentiation may be improved by providing additional growth factors and ECM components relevant for the microenvironment of mature hepatocytes^[67,68]^. In particular, mesenchymal stem cell (MSC)-conditioned medium could be utilized to more closely mimic a complex paracrine signaling^[69]^, and MSC-decellularized HFMs could serve as a source of numerous ECM components that cannot be recreated exogenously^[70]^. Low oxygen treatment is also worth considering, as drug-metabolizing enzymes are expressed in the hypoxic pericentral zone of the liver^[71]^. Moreover, longer perfusion should be applied in a millifluidic circulation (24-96 hrs) to enable detection of the metabolites of the low clearance drugs^[10,36]^. Another added value of longer perfusion may be a flow-induced shear stress that was reported to be determinant of cell maturation^[72]^. Inducing shear stress also during the cell differentiation phase may be a synergistic option to enhance the expression of drug-metabolizing enzymes. Previously, we found that this stressor improved maturation of Caco-2 cells grown on HFMs.^[14]^

We demonstrated that hepatocyte-like monolayers formed the barrier that was 10 times tighter than porous HFM walls coated with L-DOPA and laminin 332. Monolayers generated in previous studies displayed higher permeability, with only a 3-4 fold change between cell- free and cell-laden HFMs.^[14,17–19]^ To draw a reliable conclusion, permeability of differentiated Caco-2 monolayers should be examined using our system, which are known to form a tight barrier^[14]^. This will allow for determining a maximum acceptable cut-off value for permeability of hepatocyte-like monolayers. As we observed a variation in the functional barrier integrity of monolayers that we judged as confluent, establishing quantitative control criteria in addition to microscopic visualization prior to perfusion assays could reduce this variation.

We showed that our model is modular by connecting it to the kidney proximal tubule-like HFM-based model^[17]^. An in-house printed Transwell system that we previously used to establish a tissue explant-based small intestine-on-a-chip model^[37]^ may also be compatible. An ultimate goal in the future should be connecting these three models in a physiologically- relevant fashion into a multi-organ-on-a-chip platform for a full prediction of human PK/ADME profiles^[7]^. We believe that our liver-on-a-chip system can also be used for pharmacological applications other than predicting PK/ADME. In particular, the system can be maintained in differentiation medium in a broad functional time window that is suitable for examining acute hepatotoxicity of drug candidates with both an immediate and a delayed onset^[73]^, and for examining a repeated dosing^[74]^. Indeed, a recent performance assessment of 870 commercially manufactured (from *Emulate, Inc.*) human liver-like chips confirmed that they are predictive for acute drug-induced liver injury (DILI) and can be incorporated into preclinical development for hepatotoxicity testing.^[75]^ Finally, HFMs can be utilized in accordance with our protocol for creating organoid-derived perfusable monolayers from other epithelial organs, for instance, breast^[76]^, to study transepithelial transport across the highly selective blood-milk barrier to predict medication exposure via breastfeeding^[77]^. In conclusion, we provide a proof-of-concept, animal component-free, bioengineered liver organoid-on-a-chip model for examining both metabolism and transport of drugs, which is compatible with the other PK/ADME-related organ systems, namely the small-intestine and kidney.

## Author Contributions

Conceptualization: A. Myszczyszyn, B.S.; investigation: A. Myszczyszyn, A. Münch, V.L., M.B., R.S., M.K.N., T.M., D.K., S.B.; data analysis and interpretation: A. Myszczyszyn, A. Münch, T.S., F.S., M.K.N., T.M.; resources: L.L.; supervision: H.E.A., E.S., R.M., B.S.; funding acquisition: E.S., R.M., B.S.; manuscript draft preparation: A. Myszczyszyn, B.S.

## Acknowledgments

We would like to thank our consortium partners for their valuable contribution, *i.e.* Birk Poller, Markus Walles, Olivier Kretz and Katharina Root from *Novartis*; Martijn Wilmer, Ron Byron, Errol Byron and Susan Veissi from *Cell4Pharma*; Joanne Donkers from TNO; Saskia Aan and Anne Burgers from *Proefdiervrij* as well as Margot Beukers from SGF. Moreover, we acknowledge the Utrecht Sequencing Facility (USEQ) for providing sequencing service and data. USEQ is subsidized by the University Medical Center Utrecht and The Netherlands X- omics Initiative (NWO project 184.034.019).

## Data Availability Statement

The transcriptomic data were deposited to the Gene Expression Omnibus (GEO) repository (https://www.ncbi.nlm.nih.gov/geo/) with the accession number GSE269150.

---

**Fig. S1.**
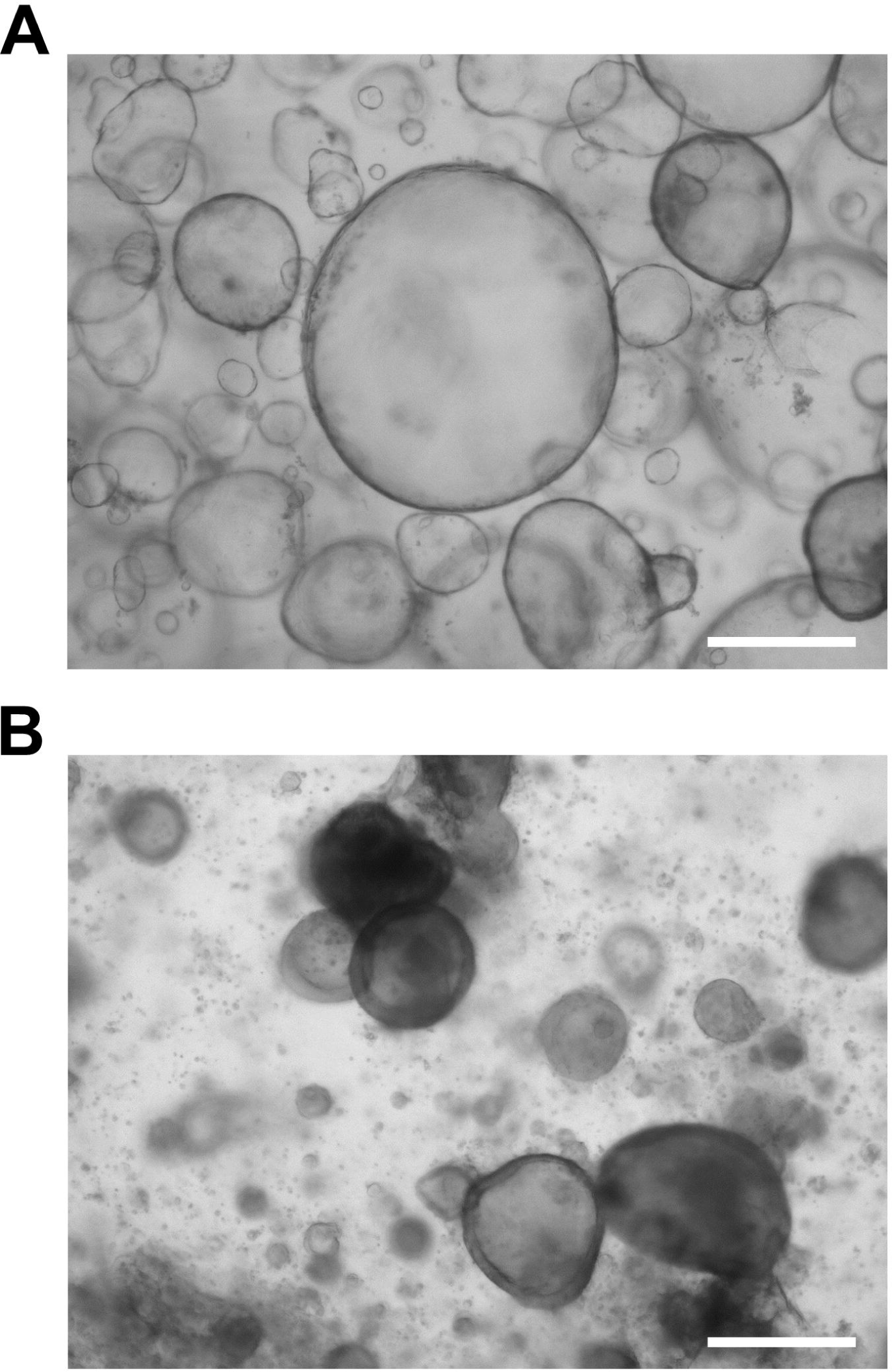
Expansion and differentiation of liver organoids in Matrigel droplets. (A) A representative brightfield image of an organoid culture after 4-7 days in EM. (B) A representative brightfield image of an expanded organoid culture after 3 days in pre-DM and 7 days in DM. Data information: scale bars, 500 µm. Four independent replicates (donors, 1-3 and 5) with multiple technical replicates (wells/droplets) were examined.

**Fig. S2.**
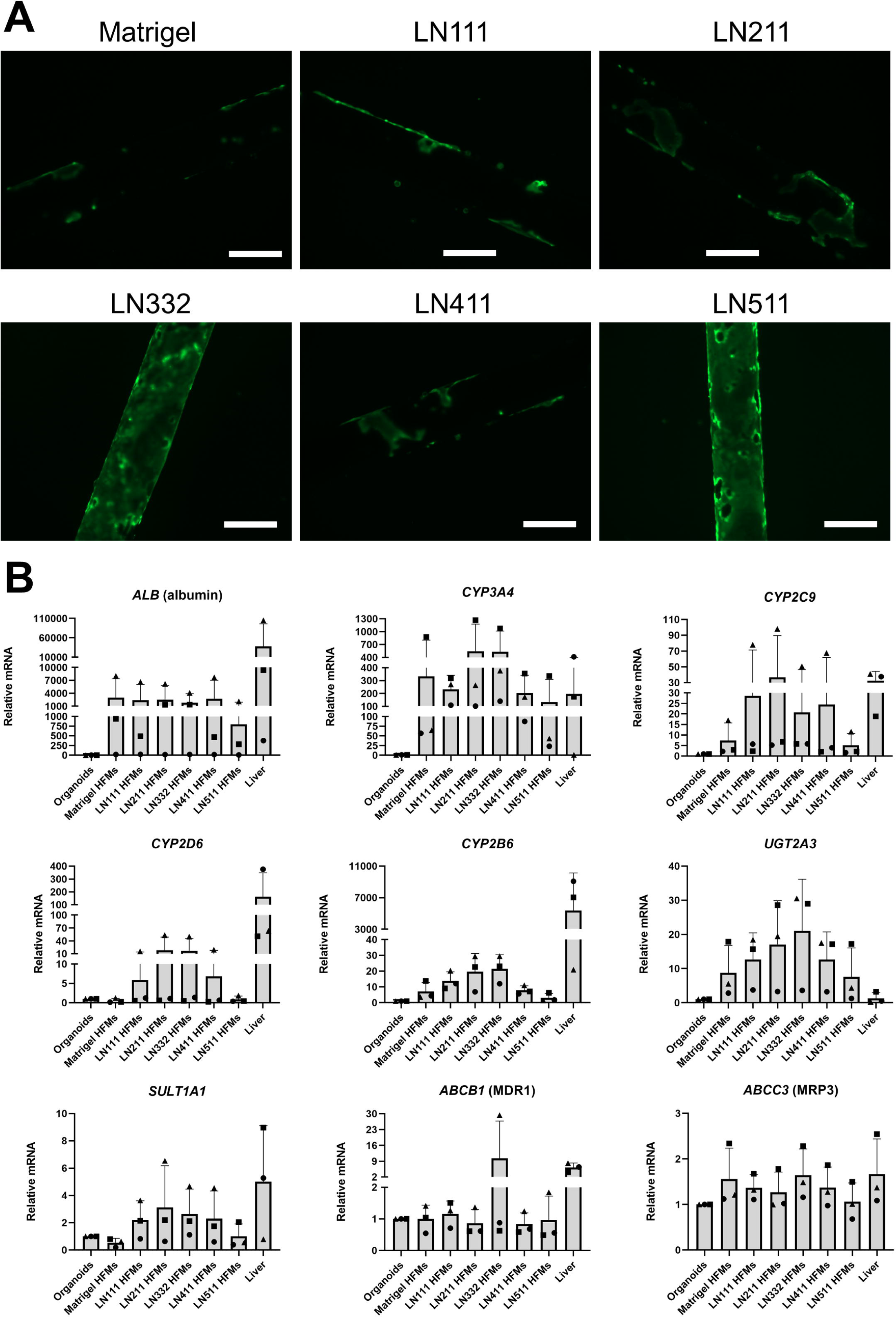
Expansion and differentiation of liver organoid fragments on statically cultured HFMs coated with L-DOPA and Matrigel, or recombinant human laminin (LN) 111, 211, 332, 411 or 511. (A) Representative images of intracellular fluorescence of calcein showing expanded monolayers after 7 days in EM. (B) Real-time qPCR analysis of the expression of *ALB* (albumin), *CYP3A4*, *CYP2C9*, *CYP2D6*, *CYP2B6*, *UGT2A3*, *SULT1A1*, *ABCB1* (MDR1) and *ABCC3* (MRP3) after 3 days in pre-DM and 7 days in DM. Data information: scale bars in A, 500 µm. Three independent replicates (donors; 2, 3, 5) with six technical replicates (monolayers) were examined. In B, the graphs show mean + SD (error bars). Three independent replicates (donors; 2, 3, 5; matched for all the conditions) in technical duplicates (wells) were examined.

**Fig. S3.**
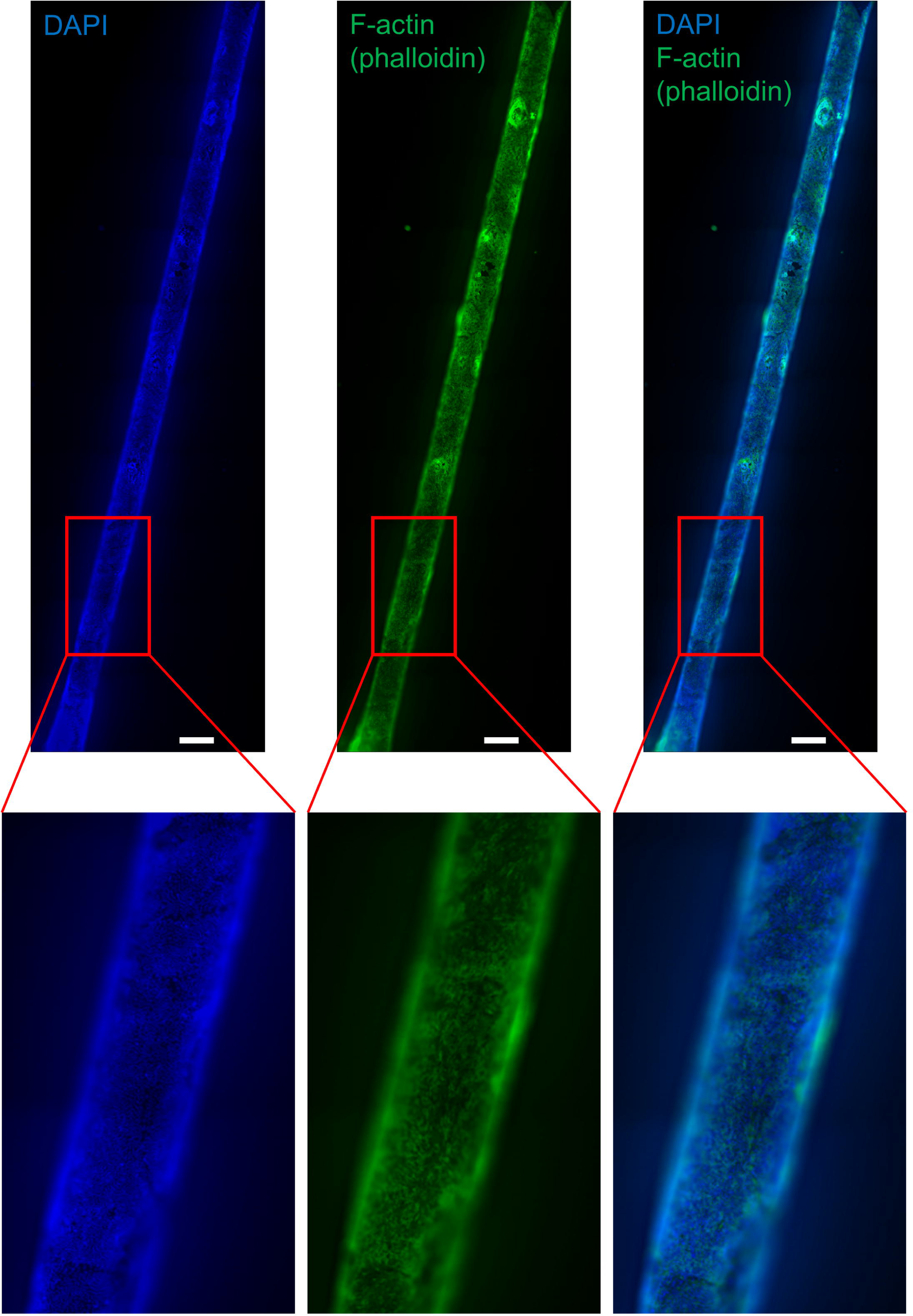
A ubiquitious expression of F-actin (phalloidin) in cell membranes of statically cultured HFM- grown monolayers. A representative whole-mount confocal immunofluorescence microscopy image of a dorsal section of a HFM. Data information: scale bars, 500 µm. Nuclei were counterstained with DAPI. Insets are enlarged in the bottom part. One independent replicate (donor, 5) in technical triplicate (monolayers) was examined.

**Fig. S4.**
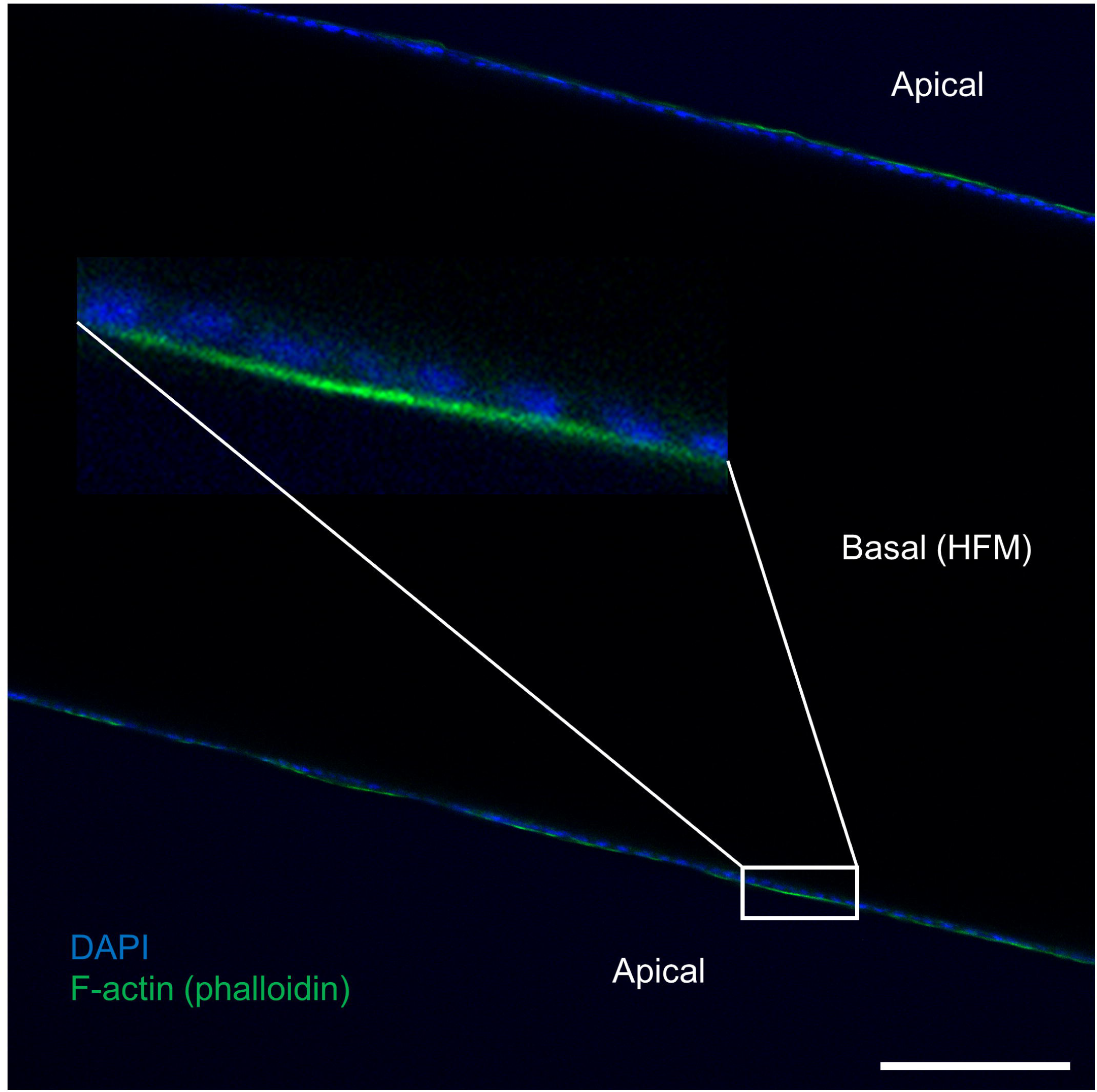
The apical localization of F-actin (phalloidin) in cell membranes of statically cultured HFM-grown monolayers. A representative whole-mount confocal immunofluorescence microscopy image of a median section of a HFM. Data information: scale bar, 100 µm. Nuclei were counterstained with DAPI. Inset is enlarged in the middle part. One independent replicate (donor, 5) in technical triplicate (monolayers) was examined.

**Fig. S5.**
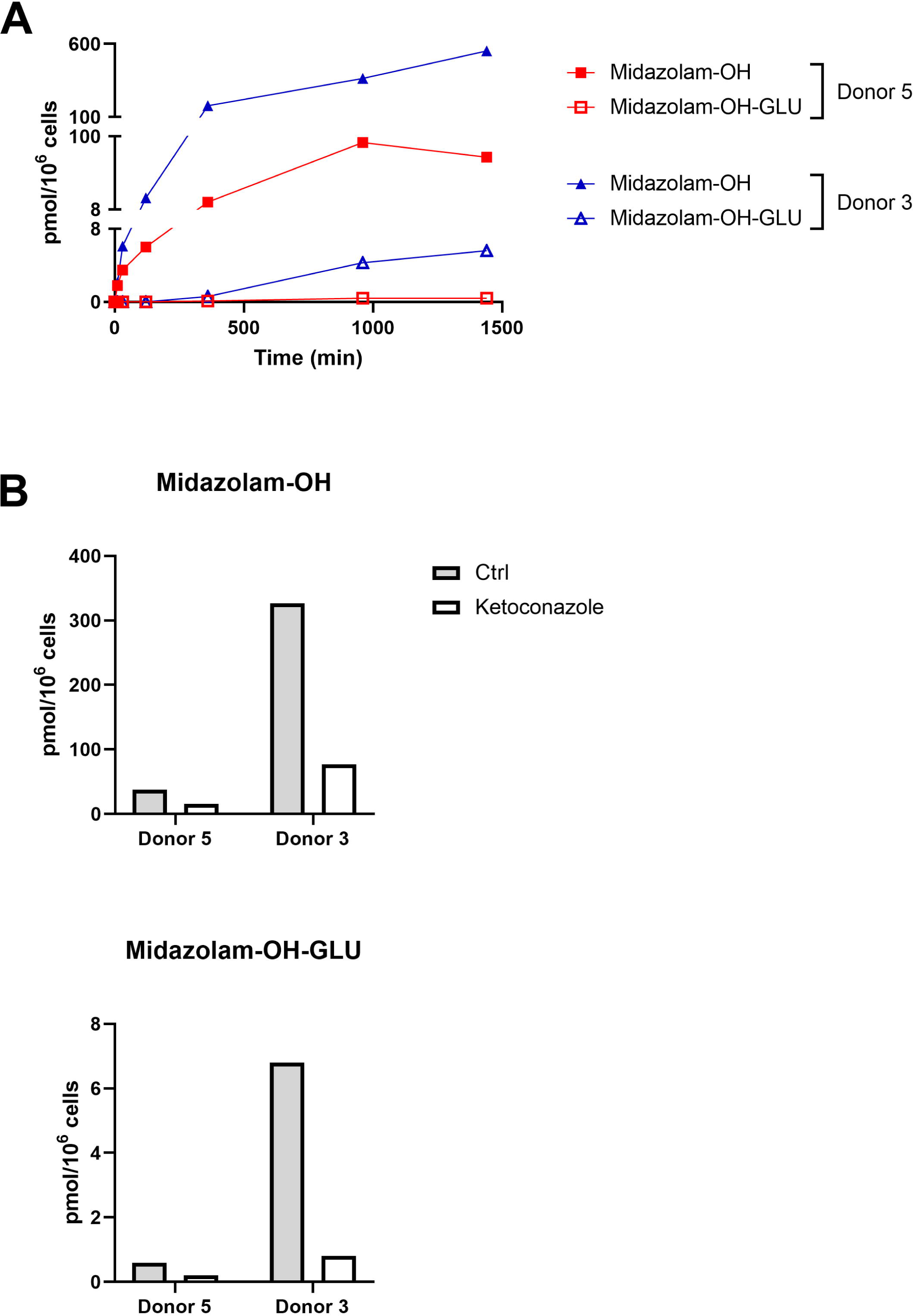
Detailed characterization of midazolam metabolism in statically cultured HFM-grown monolayers. (A) The formation of midazolam-OH and midazolam-OH-GLU over 24-h exposure with 5 µM midazolam. (B) The formation of midazolam-OH and midazolam-OH-GLU after 24-h exposure with 5 µM midazolam and treatment with ketoconazole versus EtOH (control). Data information: the curves and graphs show single technical replicates (wells/measurements, from pooled six monolayers) for two independent replicates (donors; 3, 5).

**Fig. S6.**
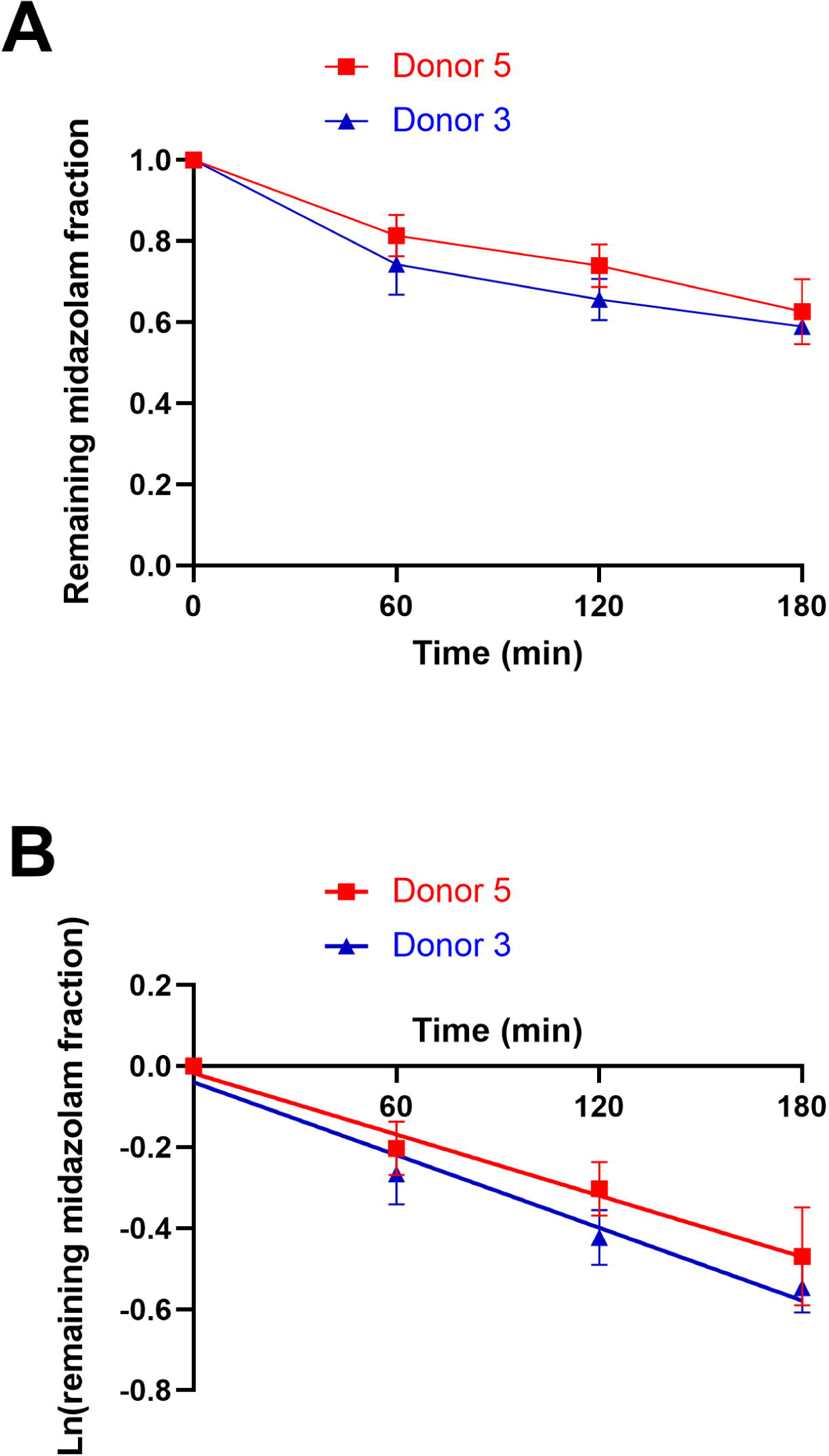
Clearance of midazolam on-a-chip. (A) A remaining fraction of 5 µM midazolam over 3-h perfusion. (B) The natural logarithm of a remaining fraction of 5 µM midazolam over 3-h perfusion. Data information: the curves show mean ± SD (error bars). Two independent replicates (donors; 3, 5) with three technical replicates (chips) were examined.

**Fig. S7.**
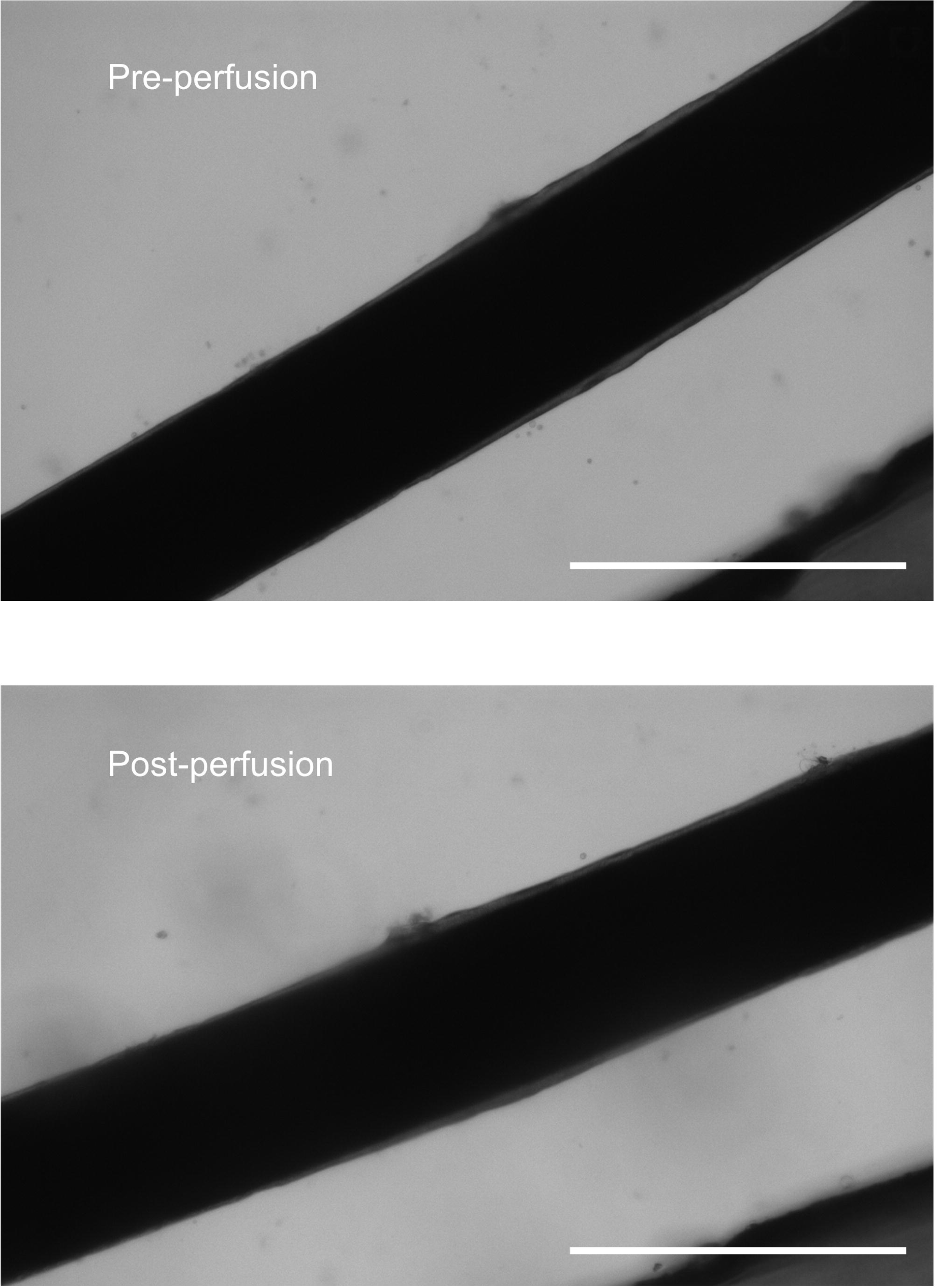
Morphology of HFM-grown monolayers before and after 3-h perfusion. Representative brightfield images of monolayers in a median section of a HFM. Data information: scale bars, 500 µm. Two independent replicates (donors; 3, 5) with two technical replicates (chips) were examined.

## References

[1] M. J. Waring, J. Arrowsmith, A. R. Leach, P. D. Leeson, S. Mandrell, R. M. Owen, G. Pairaudeau, W. D. Pennie, S. D. Pickett, J. Wang, O. Wallace, A. Weir, Nat. Rev. Drug Discov. 2015 147 2015, 14, 475.

[2] G. A. Van Norman, JACC Basic to Transl. Sci. 2019, 4, 845.

[3] D. M. Stresser, A. K. Kopec, P. Hewitt, R. N. Hardwick, T. R. Van Vleet, P. K. S. Mahalingaiah, D. O’Connell, G. J. Jenkins, R. David, J. Graham, D. Lee, J. Ekert, A. Fullerton, R. Villenave, P. Bajaj, J. R. Gosset, S. L. Ralston, M. Guha, A. Amador-Arjona, K. Khan, S. Agarwal, C. Hasselgren, X. Wang, K. Adams, G. Kaushik, A. Raczynski, K. A. Homan, Nat. Biomed. Eng. 2023 2023, 1.

[4] H. Musther, A. Olivares-Morales, O. J. D. Hatley, B. Liu, A. Rostami Hodjegan, Eur. J. Pharm. Sci. 2014, 57, 280.

[5] S. Fowler, W. L. K. Chen, D. B. Duignan, A. Gupta, N. Hariparsad, J. R. Kenny, W. G. Lai, J. Liras, J. A. Phillips, J. Gan, Lab Chip 2020, 20, 446.

[6] D. E. Ingber, Nat. Rev. Genet. 2022 238 2022, 23, 467.

[7] M. Keuper-Navis, M. Walles, B. Poller, A. Myszczyszyn, T. K. van der Made, J. Donkers, H. Eslami Amirabadi, M. J. Wilmer, S. Aan, B. Spee, R. Masereeuw, E. van de Steeg, Pharmacol. Res. 2023, 195, 106853.

[8] B. Cox, P. Barton, R. Class, H. Coxhead, C. Delatour, E. Gillent, J. Henshall, E. M. Isin, L. King, J. P. Valentin, Biomater. Biosyst. 2022, 7, 100054.

[9] K. J. Jang, M. A. Otieno, J. Ronxhi, H. K. Lim, L. Ewart, K. R. Kodella, D. B. Petropolis, G. Kulkarni, J. E. Rubins, D. Conegliano, J. Nawroth, D. Simic, W. Lam, M. Singer, E. Barale, B. Singh, M. Sonee, A. J. Streeter, C. Manthey, B. Jones, A. Srivastava, L. C. Andersson, D. Williams, H. Park, R. Barrile, J. Sliz, A. Herland, S. Haney, K. Karalis, D. E. Ingber, G. A. Hamilton, Sci. Transl. Med. 2019, 11, 5516.

[10] S. A. P. Rajan, J. Sherfey, S. Ohri, L. Nichols, J. T. Smith, P. Parekh, E. P. Kadar, F. Clark, B. T. George, L. Gregory, D. Tess, J. R. Gosset, J. Liras, E. Geishecker, R. S. Obach, M. Cirit, AAPS J. 2023, 25, 1.

[11] A. Rubiano, A. Indapurkar, R. Yokosawa, A. Miedzik, B. Rosenzweig, A. Arefin, C. M. Moulin, K. Dame, N. Hartman, D. A. Volpe, M. K. Matta, D. J. Hughes, D. G. Strauss, T. Kostrzewski, A. J. S. Ribeiro, Clin. Transl. Sci. 2021, 14, 1049.

[12] K. Sugano, M. Kansy, P. Artursson, A. Avdeef, S. Bendels, L. Di, G. F. Ecker, B. Faller, H. Fischer, G. Gerebtzoff, H. Lennernaes, F. Senner, Nat. Rev. Drug Discov. 2010 98 2010, 9, 597.

[13] V. L. Dao Thi, X. Wu, R. L. Belote, U. Andreo, C. N. Takacs, J. P. Fernandez, L. A. Vale-Silva, S. Prallet, C. C. Decker, R. M. Fu, B. Qu, K. Uryu, H. Molina, M. Saeed, E. Steinmann, S. Urban, R. R. Singaraja, W. M. Schneider, S. M. Simon, C. M. Rice, Nat. Commun. 2020, 11, 1677.

[14] P. G. M. Jochems, J. van Bergenhenegouwen, A. M. van Genderen, S. T. Eis, L. J. F. Wilod Versprille, H. J. Wichers, P. V. Jeurink, J. Garssen, R. Masereeuw, Toxicol. Vitr. 2019, 60, 1.

[15] X. Deng, G. Zhang, C. Shen, J. Yin, Q. Meng, Appl. Microbiol. Biotechnol. 2013, 97, 6943.

[16] K. C. Clause, L. J. Liu, K. Tobita, Cell Commun. Adhes. 2010, 17, 48.

[17] J. Jansen, I. E. De Napoli, M. Fedecostante, C. M. S. Schophuizen, N. V. Chevtchik, M. J. Wilmer, A. H. Van Asbeck, H. J. Croes, J. C. Pertijs, J. F. M. Wetzels, L. B. Hilbrands, L. P. Van Den Heuvel, J. G. Hoenderop, D. Stamatialis, R. Masereeuw, Sci. Reports 2015 51 2015, 5, 1.

[18] C. Chen, P. G. M. Jochems, L. Salz, K. Schneeberger, L. C. Penning, S. F. J. Van De Graaf, U. Beuers, H. Clevers, N. Geijsen, R. Masereeuw, B. Spee, Biofabrication 2018, 10, 034103.

[19] Z. Wang, J. Faria, L. J. W. van der Laan, L. C. Penning, R. Masereeuw, B. Spee, Front. Bioeng. Biotechnol. 2022, 10, DOI 10.3389/FBIOE.2022.868857/FULL.

[20] A. Marsee, F. J. M. Roos, M. M. A. Verstegen, F. Roos, M. Verstegen, H. Clevers, L. Vallier, T. Takebe, M. Huch, W. C. Peng, S. Forbes, F. Lemaigre, E. de Koning, H. Gehart, B. Spee, S. Boj, P. Baptista, K. Schneeberger, C. Soroka, M. Heim, S. Nuciforo, K. Zaret, Y. Saito, M. Lutolf, V. Cardinale, B. Simons, S. van IJzendoorn, A. Kamiya, H. Chikada, S. Wang, S. J. Mun, M. J. Son, T. T. Onder, J. Boyer, T. Sato, N. Georgakopoulos, A. Meneses, L. Broutier, L. Boulter, D. Grün, J. IJzermans, B. Artegiani, R. van Boxtel, E. Kuijk, G. Carpino, G. Peltz, J. Banales, N. Man, L. Aloia, N. LaRusso, G. George, C. Rimland, G. Yeoh, A. Grappin-Botton, D. Stange, N. Prior, J. E. E. Tirnitz-Parker, E. Andersson, C. Braconi, N. Hannan, W. Y. Lu, S. Strom, P. Sancho-Bru, S. Ogawa, V. Corbo, M. Lancaster, H. Hu, S. Fuchs, D. Hendriks, S. J. Forbes, L. J. W. van der Laan, Cell Stem Cell 2021, 28, 816.

[21] M. Huch, H. Gehart, R. Van Boxtel, K. Hamer, F. Blokzijl, M. M. A. Verstegen, E. Ellis, M. Van Wenum, S. A. Fuchs, J. De Ligt, M. Van De Wetering, N. Sasaki, S. J. Boers, H. Kemperman, J. De Jonge, J. N. M. Ijzermans, E. E. S. Nieuwenhuis, R. Hoekstra, S. Strom, R. R. G. Vries, L. J. W. Van Der Laan, E. Cuppen, H. Clevers, Cell 2015, 160, 299.

[22] M. Watanabe, H. Zemack, H. Johansson, L. Hagbard, C. Jorns, M. Li, E. Ellis, PLoS One 2016, 11, e0161383.

[23] Y. N. Kallis, A. J. Robson, J. A. Fallowfield, H. C. Thomas, M. R. Alison, N. A. Wright, R. D. Goldin, J. P. Iredale, S. J. Forbes, Gut 2011, 60, 525.

[24] K. Cameron, R. Tan, W. Schmidt-Heck, G. Campos, M. J. Lyall, Y. Wang, B. Lucendo-Villarin, D. Szkolnicka, N. Bates, S. J. Kimber, J. G. Hengstler, P. Godoy, S. J. Forbes, D. C. Hay, Stem Cell Reports 2015, 5, 1250.

[25] D. F. B. Malta, N. E. Reticker-Flynn, C. L. Da Silva, J. M. S. Cabral, H. E. Fleming, K. S. Zaret, S. N. Bhatia, G. H. Underhill, Acta Biomater. 2016, 34, 30.

[26] L. K. Kanninen, R. Harjumäki, P. Peltoniemi, M. S. Bogacheva, T. Salmi, P. Porola, J. Niklander, T. Smutný, A. Urtti, M. L. Yliperttula, Y. R. Lou, Biomaterials 2016, 103, 86.

[27] J. Ong, M. P. Serra, J. Segal, A. M. Cujba, S. S. Ng, R. Butler, V. Millar, S. Hatch, S. Zimri, H. Koike, K. Chan, A. Bonham, M. Walk, T. Voss, N. Heaton, R. Mitry, A. Dhawan, D. Ebner, D. Danovi, H. Nakauchi, S. T. Rashid, Stem Cell Reports 2018, 10, 693.

[28] K. Takayama, Y. Morisaki, S. Kuno, Y. Nagamoto, K. Harada, N. Furukawa, M. Ohtaka, K. Nishimura, K. Imagawa, F. Sakurai, M. Tachibana, R. Sumazaki, E. Noguchi, M. Nakanishi, K. Hirata, K. Kawabata, H. Mizuguchi, Proc. Natl. Acad. Sci. U. S. A. 2014, 111, 16772.

[29] Z. Wang, S. Ye, L. J. W. van der Laan, K. Schneeberger, R. Masereeuw, B. Spee, Adv. Healthc. Mater. 2024, DOI 10.1002/ADHM.202401511.

[30] S. Ye, J. W. B. Boeter, M. Mihajlovic, F. G. van Steenbeek, M. E. van Wolferen, L. A. Oosterhoff, A. Marsee, M. Caiazzo, L. J. W. van der Laan, L. C. Penning, T. Vermonden, B. Spee, K. Schneeberger, Adv. Funct. Mater. 2020, 30, DOI 10.1002/ADFM.202000893.

[31] M. C. Bouwmeester, Y. Tao, S. Proença, F. G. van Steenbeek, R. A. Samsom, S. M. Nijmeijer, T. Sinnige, L. J. W. van der Laan, J. Legler, K. Schneeberger, N. I. Kramer, B. Spee, Molecules 2023, 28, DOI 10.3390/MOLECULES28020621/S1.

[32] Y. Zhang, G. Parmigiani, W. E. Johnson, NAR Genomics Bioinforma. 2020, 2, DOI 10.1093/NARGAB/LQAA078.

[33] M. D. Robinson, A. Oshlack, Genome Biol. 2010, 11, 1.

[34] M. D. Robinson, D. J. McCarthy, G. K. Smyth, Bioinformatics 2010, 26, 139.

[35] H. Wickham, Ggplot2: Elegant Graphics for Data Analysis. Second Edition. Springer., 2016.

[36] L. Docci, N. Milani, T. Ramp, A. A. Romeo, P. Godoy, D. O. Franyuti, S. Krähenbühl, M. Gertz, A. Galetin, N. Parrott, S. Fowler, Lab Chip 2022, 22, 1187.

[37] H. Eslami Amirabadi, J. M. Donkers, E. Wierenga, B. Ingenhut, L. Pieters, L. Stevens, T. Donkers, J. Westerhout, R. Masereeuw, I. Bobeldijk-Pastorova, I. Nooijen, E. Van De Steeg, Lab Chip 2022, 22, 326.

[38] T. Yamashita, T. Inui, J. Yokota, K. Kawakami, G. Morinaga, M. Takatani, D. Hirayama, R. Nomoto, K. Ito, Y. Cui, S. Ruez, K. Harada, W. Kishimoto, H. Nakase, H. Mizuguchi, Mol. Ther. Methods Clin. Dev. 2021, 22, 263.

[39] A. Domogatskaya, S. Rodin, K. Tryggvason, Annu. Rev. Cell Dev. Biol. 2012, 28, 523.

[40] A. Galetin, K. L. R. Brouwer, D. Tweedie, K. Yoshida, N. Sjöstedt, L. Aleksunes, X. Chu, R. Evers, M. J. Hafey, Y. Lai, P. Matsson, A. Riselli, H. Shen, A. Sparreboom, M. V. S. Varma, J. Yang, X. Yang, S. W. Yee, M. J. Zamek-Gliszczynski, L. Zhang, K. M. Giacomini, Nat. Rev. Drug Discov. 2024 234 2024, 23, 255.

[41] A. Saravanakumar, A. Sadighi, R. Ryu, F. Akhlaghi, Clin. Pharmacokinet. 2019, 58, 1281.

[42] T. Fukami, T. Yokoi, M. Nakajima, Annu. Rev. Pharmacol. Toxicol. 2022, 62, 405.

[43] A. Rowland, J. O. Miners, P. I. Mackenzie, Int. J. Biochem. Cell Biol. 2013, 45, 1121.

[44] A. Isvoran, Y. Peng, S. Ceauranu, L. Schmidt, A. B. Nicot, M. A. Miteva, Drug Discov. Today 2022, 27, 103349.

[45] J. D. Hayes, J. U. Flanagan, I. R. Jowsey, Annu. Rev. Pharmacol. Toxicol. 2005, 45, 51.

[46] P. Caetano-Pinto, P. Nordell, T. Nieskens, K. Haughan, K. S. Fenner, S. H. Stahl, ALTEX - Altern. to Anim. Exp. 2023, 40, 408.

[47] A. M. van Genderen, K. Jansen, M. Kristen, J. van Duijn, Y. Li, C. C. L. Schuurmans, J. Malda, T. Vermonden, J. Jansen, R. Masereeuw, M. Castilho, Front. Bioeng. Biotechnol. 2021, DOI 10.3389/fbioe.2020.617364.

[48] D. Spaggiari, L. Geiser, Y. Daali, S. Rudaz, J. Pharm. Biomed. Anal. 2014, 101, 221.

[49] R. O. Juvonen, A. T. Heikkinen, O. Kärkkäinen, R. Jehangir, J. Huuskonen, J. Troberg, H. Raunio, O. T. Pentikäinen, M. Finel, Eur. J. Pharm. Sci. 2020, 141, 105118.

[50] C. Y. J. Hsieh, M. Sun, G. Osborne, K. Ricker, F. C. Tsai, K. Li, R. Tomar, J. Phuong, R. Schmitz, M. S. Sandy, Int. J. Toxicol. 2019, 38, 501.

[51] M. T. Donato, D. Hallifax, L. Picazo, J. V. Castell, J. B. Houston, M. J. Gomez-Lechón, A. Lahoz, Drug Metab. Dispos. 2010, 38, 1449.

[52] N. A. Kratochwil, C. Meille, S. Fowler, F. Klammers, A. Ekiciler, B. Molitor, S. Simon, I. Walter, C. McGinnis, J. Walther, B. Leonard, M. Triyatni, H. Javanbakht, C. Funk, F. Schuler, T. Lavé, N. J. Parrott, AAPS J. 2017, 19, 534.

[53] B. Zhu, D. Bush, G. A. Doss, S. Vincent, R. B. Franklin, S. Xu, Drug Metab. Dispos. 2008, 36, 331.

[54] D. J. Greenblatt, Y. Zhao, K. Venkatakrishnan, S. X. Duan, J. S. Harmatz, S. J. Parent, M. H. Court, L. L. Von Moltke, J. Pharm. Pharmacol. 2011, DOI 10.1111/j.2042-7158.2010.01202.x.

[55] E. Järvinen, F. Deng, W. Kiander, A. Sinokki, H. Kidron, N. Sjöstedt, Front. Pharmacol. 2022, 12, DOI 10.3389/FPHAR.2021.802539/FULL.

[56] T. Bricks, J. Hamon, M. J. Fleury, R. Jellali, F. Merlier, Y. E. Herpe, A. Seyer, J. M. Regimbeau, F. Bois, E. Leclerc, Biopharm. Drug Dispos. 2015, 36, 275.

[57] N. Milani, N. Parrott, D. Ortiz Franyuti, P. Godoy, A. Galetin, M. Gertz, S. Fowler, Lab Chip 2022, 22, 2853.

[58] J. M. Prot, L. Maciel, T. Bricks, F. Merlier, J. Cotton, P. Paullier, F. Y. Bois, E. Leclerc, Biotechnol. Bioeng. 2014, 111, 2027.

[59] N. Tsamandouras, W. L. K. Chen, C. D. Edington, C. L. Stokes, L. G. Griffith, M. Cirit, AAPS J. 2017, 19, 1499.

[60] G. G. Graham, J. Punt, M. Arora, R. O. Day, M. P. Doogue, J. K. Duong, T. J. Furlong, J. R. Greenfield, L. C. Greenup, C. M. Kirkpatrick, J. E. Ray, P. Timmins, K. M. Williams, Clin. Pharmacokinet. 2011, 50, 81.

[61] Y. Zhao, S. Landau, S. Okhovatian, C. Liu, R. X. Z. Lu, B. F. L. Lai, Q. Wu, J. Kieda, K. Cheung, S. Rajasekar, K. Jozani, B. Zhang, M. Radisic, Nat. Rev. Bioeng. 2024 2024, 1.

[62] S. Lorenzini, T. G. Bird, L. Boulter, C. Bellamy, K. Samuel, R. Aucott, E. Clayton, P. Andreone, M. Bernardi, M. Golding, M. R. Alison, J. P. Iredale, S. J. Forbes, Gut 2010, 59, 645.

[63] F. Schutgens, H. Clevers, Annu. Rev. Pathol. Mech. Dis. 2020, 15, 211.

[64] M. Yamada, H. Okada, Y. Kikkawa, A. Miyajima, T. Itoh, Biochem. Biophys. Res. Commun. 2020, 524, 465.

[65] Y. Zhou, V. M. Lauschke, Pharmacogenomics J. 2022 225 2022, 22, 284.

[66] L. Kuepfer, C. Niederalt, T. Wendl, J. F. Schlender, S. Willmann, J. Lippert, M. Block, T. Eissing, D. Teutonico, CPT Pharmacometrics Syst. Pharmacol. 2016, 5, 516.

[67] C. Chen, A. Soto-Gutierrez, P. M. Baptista, B. Spee, Gastroenterology 2018, 154, 1258.

[68] D. Hendriks, B. Artegiani, T. Margaritis, I. Zoutendijk, S. Chuva de Sousa Lopes, H. Clevers, Nat. Commun. 2024 151 2024, 15, 1.

[69] T. Takebe, R. R. Zhang, H. Koike, M. Kimura, E. Yoshizawa, M. Enomura, N. Koike, K. Sekine, H. Taniguchi, Nat. Protoc. 2014 92 2014, 9, 396.

[70] X. Zhang, X. Chen, H. Hong, R. Hu, J. Liu, C. Liu, Bioact. Mater. 2022, 10, 15.

[71] J. Paris, N. C. Henderson, Hepatology 2022, 76, 1219.

[72] M. Hofer, M. P. Lutolf, Nat. Rev. Mater. 2021, 6, 402.

[73] F. Bessone, N. Hernandez, M. Tagle, M. Arrese, R. Parana, N. Mendez-Sánchez, E. Ridruejo, M. Mendizabal, L. Dagher, F. Contreras, E. Fassio, M. Pesoa, J. Brahm, M. Silva, Ann. Hepatol. 2021, 24, 100321.

[74] I. Maschmeyer, T. Hasenberg, A. Jaenicke, M. Lindner, A. K. Lorenz, J. Zech, L. A. Garbe, F. Sonntag, P. Hayden, S. Ayehunie, R. Lauster, U. Marx, E. M. Materne, Eur. J. Pharm. Biopharm. 2015, 95, 77.

[75] L. Ewart, A. Apostolou, S. A. Briggs, C. V Carman, J. T. Chaff, A. R. Heng, S. Jadalannagari, J. Janardhanan, K.-J. Jang, S. R. Joshipura, M. M. Kadam, M. Kanellias, V. J. Kujala, G. Kulkarni, C. Y. Le, C. Lucchesi, D. V Manatakis, K. K. Maniar, M. E. Quinn, J. S. Ravan, A. C. Rizos, J. F. K. Sauld, J. D. Sliz, W. Tien-Street, D. Ramos Trinidad, J. Velez, M. Wendell, O. Irrechukwu, P. K. Mahalingaiah, D. E. Ingber, J. W. Scannell, D. Levner, H. John, Commun. Med. 2022 21 2022, 2, 1.

[76] N. Sachs, J. de Ligt, O. Kopper, E. Gogola, G. Bounova, F. Weeber, A. V. Balgobind, K. Wind, A. Gracanin, H. Begthel, J. Korving, R. van Boxtel, A. A. Duarte, D. Lelieveld, A. van Hoeck, R. F. Ernst, F. Blokzijl, I. J. Nijman, M. Hoogstraat, M. van de Ven, D. A. Egan, V. Zinzalla, J. Moll, S. F. Boj, E. E. Voest, L. Wessels, P. J. van Diest, S. Rottenberg, R. G. J. Vries, E. Cuppen, H. Clevers, Cell 2018, DOI 10.1016/j.cell.2017.11.010.

[77] N. Nauwelaerts, N. Deferm, A. Smits, C. Bernardini, B. Lammens, P. Gandia, A. Panchaud, H. Nordeng, M. L. Bacci, M. Forni, D. Ventrella, K. Van Calsteren, A. DeLise, I. Huys, M. Bouisset-Leonard, K. Allegaert, P. Annaert, Biomed. Pharmacother. 2021, 136, 111038.

